# A hidden gene in astroviruses encodes a cell-permeabilizing protein involved in virus release

**DOI:** 10.1101/661579

**Authors:** Valeria Lulla, Andrew E. Firth

## Abstract

Human astroviruses are small nonenveloped viruses with positive-sense single-stranded RNA genomes that contain three main open reading frames: ORF1a, ORF1b and ORF2. Astroviruses cause acute gastroenteritis in children worldwide and have been associated with encephalitis and meningitis in immunocompromised individuals. Through comparative genomic analysis of >400 astrovirus sequences, we identified a conserved “ORFX” overlapping the capsid-encoding ORF2 in genogroup I, III and IV astroviruses. ORFX appears to be subject to purifying selection, consistent with it encoding a functional protein product, termed XP. Using ribosome profiling of cells infected with human astrovirus 1, we confirm initiation at the ORFX AUG. XP-knockout astroviruses are strongly attenuated and after passaging can partly restore viral titer via pseudo-reversions, thus demonstrating that XP plays an important role in virus growth. To further investigate XP, we developed an astrovirus replicon system. We demonstrate that XP has only minor effects on RNA replication and structural protein production. Instead, XP associates with the plasma membrane with an extracellular N-terminus topology and promotes efficient virus release. Using two different assays, we show that expression of human or related astrovirus XPs leads to cell permeabilization, suggesting a viroporin-like activity. The discovery of XP advances our knowledge of these important human viruses and opens a new direction of research into astrovirus replication and pathogenesis.

## INTRODUCTION

Humans astroviruses (HAstVs) belong to genus *Mamastrovirus* within the family *Astroviridae*. They are considered to be the second most common cause of gastroenteritis in infants after rotavirus ^1^. Recently, and especially in immunocompromised individuals, astroviruses have also been associated with extra-intestinal infections including fatal meningitis and encephalitis ^2^. Despite their undoubted importance, astroviruses still represent one of the least studied groups of human positive-sense RNA viruses. The genome contains three main open reading frames: ORF1a encoding a nonstructural polyprotein (nsP1a), ORF1b encoding the RNA-dependent RNA polymerase (RdRp), and ORF2 encoding the capsid protein (CP) (Fig. 1A) ^3^. ORF1a and ORF1b are translated from the genomic RNA (gRNA), with expression of ORF1b depending on programmed ribosomal frameshifting whereby a proportion of ribosomes translating ORF1a make a −1 nucleotide shift into ORF1b. Frameshifting occurs at a conserved A_AAA_AAC sequence within the ORF1a/ORF1b overlap region and depends on a 3′-adjacent stimulatory RNA stem-loop structure ^4^. ORF2 is translated from a subgenomic RNA (sgRNA) that is produced during virus infection.

**Figure 1.**
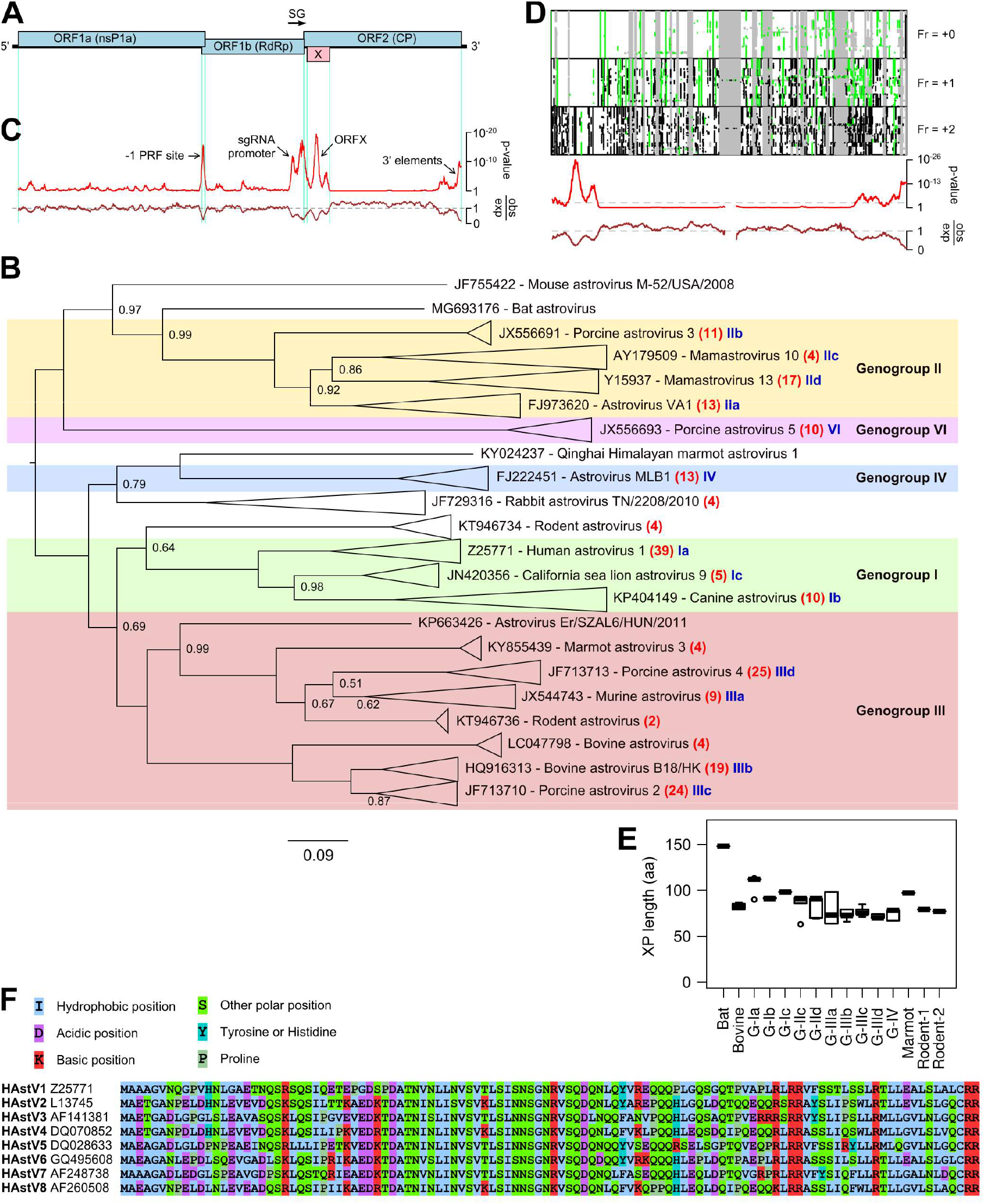
Comparative genomic analysis of astroviruses. **(A)** Map of the human astrovirus genome showing the three main ORFs (blue) and the overlapping ORFX (pink). **(B)** Phylogenetic tree of mammalian astroviruses. The tree, calculated with MrBayes, is based on ORF1b amino acid sequences obtained from 221 full-length genomes. Related groups of sequences (indicated by isosceles triangles) have been replaced in the figure by a single representative accession number and virus name; the total number of sequences in each group is shown in red (see Fig. S1 for the complete tree). Genogroups (according to Yokoyama et al.^8^) are indicated by background shading. Subgroups of sequences, defined for the purposes of this study only, are indicated in blue (Ia, Ib, etc). The tree is midpoint rooted and nodes are labelled with posterior probability values if different from 1.00. **(C)** Analysis of conservation at ORF1a-ORF1b-ORF2 synonymous sites. The red line shows the probability that the observed conservation could occur under a null model of neutral evolution at synonymous sites, whereas the brown line depicts the ratio of the observed number of substitutions to the number expected under the null model. Inferred elements corresponding to regions of enhanced synonymous site conservation are indicated. **(D)** Analysis of an alignment of 127 human and 5 feline astrovirus ORF2 sequences. The upper three panels show the positions of alignment gaps (grey), stop codons (black) and AUG codons (green) in each reading frame. Below, is shown the analysis of conservation at synonymous sites. **(E)** Box plots of XP length for different astrovirus clades: centre lines = medians; boxes = interquartile ranges; whiskers extend to most extreme data point within 1.5 × interquartile range from the box; circles = outliers; *n* = number of sequences as shown in B. **(F)** Alignment of XP sequences from representative HAstVs.

Previously, using comparative genomics we identified a conserved fourth ORF (ORFX) in genogroup I astroviruses, which appears to be subject to purifying selection and therefore is likely to encode a functional protein product, termed the X protein (XP, 12 kDa, 112 aa) ^5^. ORFX overlaps the 5′ region of the capsid-encoding ORF2 and is predicted to be translatable via ribosomal leaky scanning, which is expected to be enhanced by the very short leader on the sgRNA ^3,6,7^. However, despite the comparative genomic predictions, the existence, relevance, and function of XP have never been demonstrated. Here we perform a functional molecular dissection of ORFX, providing insight into the simultaneous orchestration of several overlapping functional elements in a small human-pathogenic RNA virus. We show that ORFX is translated during virus infection, XP knockout mutant viruses are highly attenuated, and XP is a transmembrane (TM) protein that promotes efficient virus release via a viroporin-like activity. Through an extended bioinformatic analysis, we predict that related XP proteins are widely encoded in other mammalian astroviruses including the divergent “MLB” group of human-infecting astroviruses.

## RESULTS

### Comparative genomic analysis reveals a potential ORFX in genogroup I, III and IV astroviruses

Since there are many more astrovirus sequences available now than in 2010, we began by repeating our previous comparative genomic analysis but this time also extending to other astrovirus genogroups. All mammalian astrovirus complete or nearly complete genome sequences were obtained from the National Center for Biotechnology Information (NCBI), ORF1b (RdRp) amino acid sequences were extracted and aligned, and a phylogenetic tree constructed (Fig. S1). We follow the genogroups defined in Yokoyama et al. ^8^. Although the more-divergent CP sequences may provide a less robust phylogenetic analysis, we also constructed a CP-based phylogenetic tree to test for possible recombination (Fig. S2). At the level of genogroup the ORF1b and CP trees were consistent, although within genogroups there were some differences between the topologies of the two trees. For simplicity, we used the ORF1b tree to define astrovirus clades for full-genome analyses.

We grouped astrovirus sequences into clades (Fig. 1B), generated codon-based alignments of concatenated ORF1a-ORF1b-ORF2 coding sequences within each clade, and analyzed the alignments with synplot2 which tests for regions where synonymous substitutions occur less often than average for the sequence alignment ^9^. Regions with significantly enhanced synonymous site conservation typically harbour overlapping functional elements which constraint sequence evolution ^9^. The analysis of subgroup “Ia” astroviruses (Fig. 1B; classical HAstVs besides some feline and sea lion astroviruses) revealed conserved elements at the junction of ORF1a and ORF1b (representing the ribosomal frameshifting signal and ORF1a/ORF1b overlap), upstream of ORF2 (presumed to represent elements involved in sgRNA synthesis), towards the 3′ end of ORF2 (possible replication elements), and overlapping the 5′ end of ORF2 (the putative overlapping ORFX) (Fig. 1C). We applied the same analysis to 13 other astrovirus genogroup or sub-genogroup clades, revealing conserved regions overlapping the 5′ end of ORF2 in genogroup I, III and IV astroviruses but, generally (see below for exceptions), not in genogroup II or VI astroviruses (Fig. S3).

We repeated the synonymous site conservation analysis using clades, sequence alignments and phylogenetic trees based on only ORF2 in order to rule out potential artefacts as a result of possible recombination between the nonstructural and structural modules of the virus genome, and to include additional part-genome sequences (total 415 sequences with ORF2 coverage; Fig. 1D; Fig. S4). This analysis revealed conserved regions overlapping the 5′ end of ORF2 throughout genogroup I, III and IV astroviruses but, generally, not in genogroup II or VI astroviruses (Fig. S5). Those clades with synonymous site conservation also contain a conserved overlapping +1 frame ORF, whereas those sequences without synonymous site conservation generally also lacked a conserved overlapping +1 frame ORF (Fig. S5). Interestingly, we also detected synonymous site conservation in two subgroups of genogroup II astroviruses (Fig. S3G; Fig. S3H). In subgroup “IId”, the conservation coincided with a conserved overlapping +1 frame ORF (Fig. S5E); however in subgroup “IIc”, there was no conserved ORF in the +1 reading frame. Instead, we observed an ORF in the −1 reading frame with no AUG codon (ORFY, Fig. S5D) but with conserved signals for −1 programmed ribosomal frameshifting ^10^, namely a conserved A_AAA_AAZ (Z = A, C or U) slippery heptanucleotide and a 3′-adjacent RNA stem-loop (or pseudoknot structure) (Fig. S6). It seems plausible that ORFX and ORFY in genogroup II astroviruses may have evolved independently of ORFX in other astrovirus genogroups (see Discussion).

Evidence for ORFX was also found in a clade of rodent astroviruses of unassigned genogroup (Fig. S3D; Fig. S5L). Further, a number of unclustered divergent sequences also contain long +1 frame ORFs overlapping the 5′ end of ORF2 that potentially encode XP proteins (Fig. S7). Note that genogroup V – represented by a single partial sequence, FJ890355 (bottlenose dolphin astrovirus 1) – is one of these.

The putative XP proteins encoded by genogroup I, III and IV astroviruses typically range from 70–112 aa in length and 7.5–12.3 kDa in molecular mass (Fig. 1E-F; Table S1). Similar to other overlapping genes, which generally evolve *de novo* ^11^, the XP peptide sequences show little to no homology to known protein domains. Many XP sequences contain a predicted TM domain, and some other XPs contain a stretch of hydrophobic amino acids that resembles a TM domain despite being scored below threshold by Phobius (Fig. S8, Fig. S9).

### Ribosome profiling confirms ORFX initiation in HAstV1-infected cells

To test for XP expression in infected cells, we first raised various antibodies against XP peptides and we also tested the viability of tagged-XP viruses. However, neither antibody nor tagged virus approaches were successful. Thus we turned to ribosome profiling (Ribo-Seq). Ribo-Seq is a high throughput sequencing technique that globally maps the footprints of initiating or elongating 80S ribosomes on mRNAs ^12,13^. We infected Caco2 cells and performed Ribo-Seq at 12 hours post infection (hpi). Ribo-Seq quality was assessed as previously described ^14^ (Fig. S10). Using flash-freezing with no drug pre-treatment (NT), we mapped the translational landscape of the HAstV1 genome (Fig. 2A). ORF2 is translated at ~9× the level of ORF1a, whereas ORF1b is translated at ~25% the level of ORF1a, indicating a ribosomal frameshifting efficiency of ~25%. The higher expression of ORF2 is likely a result of higher levels of sgRNA than gRNA in the translation pool. To identify translation initiation sites, we utilized the translation inhibitor lactimidomycin (LTM), which acts preferentially on the initiating ribosome but not on the elongating ribosome ^15^. LTM binds to the 80S ribosome already assembled at the initiation codon and occupies the empty exit (E-site) of initiating ribosomes, thus completely blocking translocation. Using this approach, we confirm the two previously known initiation sites, i.e. for ORF1a and ORF2, and also identify substantial initiation at the ORFX start codon (Fig. 2B), this being the third largest peak in the LTM virus profile.

**Figure 2.**
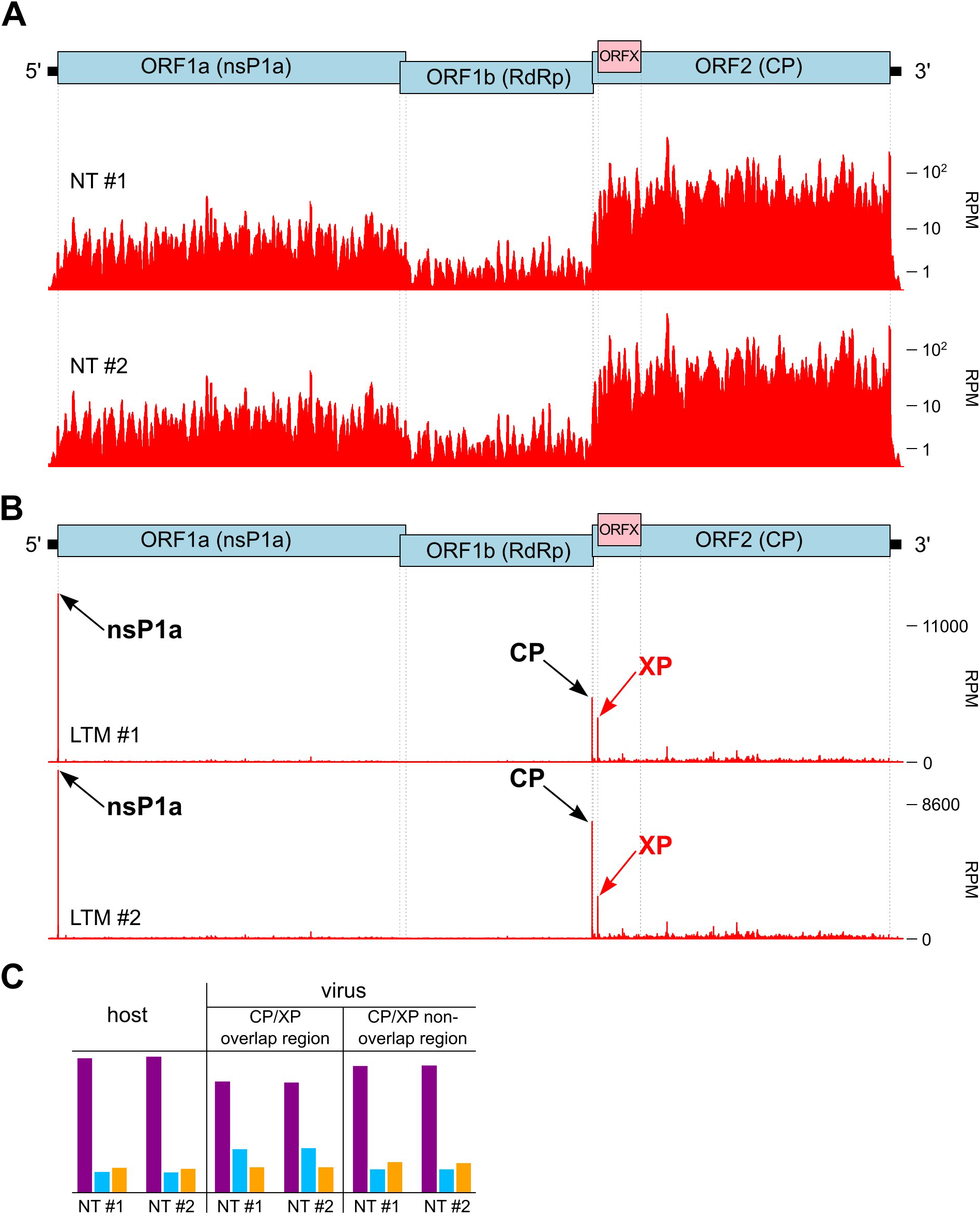
Ribosome profiling of astrovirus-infected cells. Cells were harvested at 12 hpi and either flash frozen with no pre-treatment (NT), or pre-treated with lactimidomycin for 30 minutes followed by flash freezing (LTM). **(A)** RPF densities in reads per million mapped reads (RPM) for NT repeats, smoothed with a 15-nt sliding window. **(B)** RPF densities in reads per million mapped reads (RPM) for LTM repeats, at single-nucleotide resolution. **(C)** For NT samples, phasing of 5′ ends of RPFs that map to host coding sequences, the part of ORF2 that is overlapped by ORFX, and the part of ORF2 that is not overlapped by ORFX.

It is worth noting that the short leader of the sgRNA may result in protection by the ribosome of the 5′ end of ORF2 initiation footprints from the RNase I nuclease, so that many or most ORF2 initiation footprints might retain the viral VPg protein that is thought to be covalently linked to the 5′ end of genomic and subgenomic RNAs. Such reads will not ligate to the adapter oligonucleotides and thus will be excluded from sequencing. This may explain why the ORF2 initiation peak is smaller than the ORF1a initiation peak, even though ORF2 is expressed at much higher levels than ORF1a. Thus we cannot quantify the ORF2:ORFX expression ratio from the LTM data.

Ribosome profiling of eukaryotic systems typically has the characteristic that mappings of the 5′ end positions of ribosome protected fragments (RPFs) to coding sequences reflect the triplet periodicity (herein referred to as “phasing”) of genetic decoding. For our datasets, the great majority of RPF 5′ ends map to the first nucleotide of codons (Fig. 2C, left). Reads mapping to the ORF2/ORFX part of the genome (NT samples) were quantified in the three possible phases. In the region of ORF2 that is overlapped by ORFX we observed an increased number of reads mapping in the +1 phase relative to the ORF2 reading frame (Fig. 2C, middle) compared to the region of ORF2 that is not overlapped by ORFX (Fig. 2C, right) or the coding regions of host mRNAs (Fig. 2C, left). Quantification of the differences in phasing indicated that ORFX is translated at ~27% of the level of ORF2, although it should be noted that reporter assays (see below) are expected to provide more accurate quantification than RPF phasing analysis.

### XP-knockout viruses are attenuated but pseudo-revert on passage

To evaluate the significance of ORFX in the context of virus infection, a set of mutant virus genomes was created based on the pAVIC1 infectious clone ^16^ by introducing mutations that knock out ORFX without affecting the CP amino acid sequence (Fig. 3A). Four independent mutations were introduced to guard against the possibility of affecting potential RNA secondary structures overlapping with this region, resulting in pAVIC1-AUGm (AUG to ACG), pAVIC1-PTC1 (stop codon after 20 amino acids), pAVIC1-2×PTC (double stop codon after 20 amino acids) and pAVIC1-PTC2 (stop codon after 73 amino acids) (Fig. 3B; Fig. S11). Corresponding T7 RNA transcripts were used for virus rescue in Huh7.5.1 cells followed by infection in Caco2 cells. All four ORFX knockout viruses were strongly attenuated compared to wt virus (Fig. 3C). After seven blind passages in Caco2 cells, two mutant viruses (pAVIC1-AUGm and pAVIC1-PTC1) demonstrated an ~1 log increase in virus titers, which was achieved via a pseudo-reversion (pAVIC1-AUGm) or 5 or 8 codon deletions (pAVIC1-PTC1) (Fig. 3C). All three mutations would result in restoration of XP expression, confirming its importance in virus growth.

**Figure 3.**
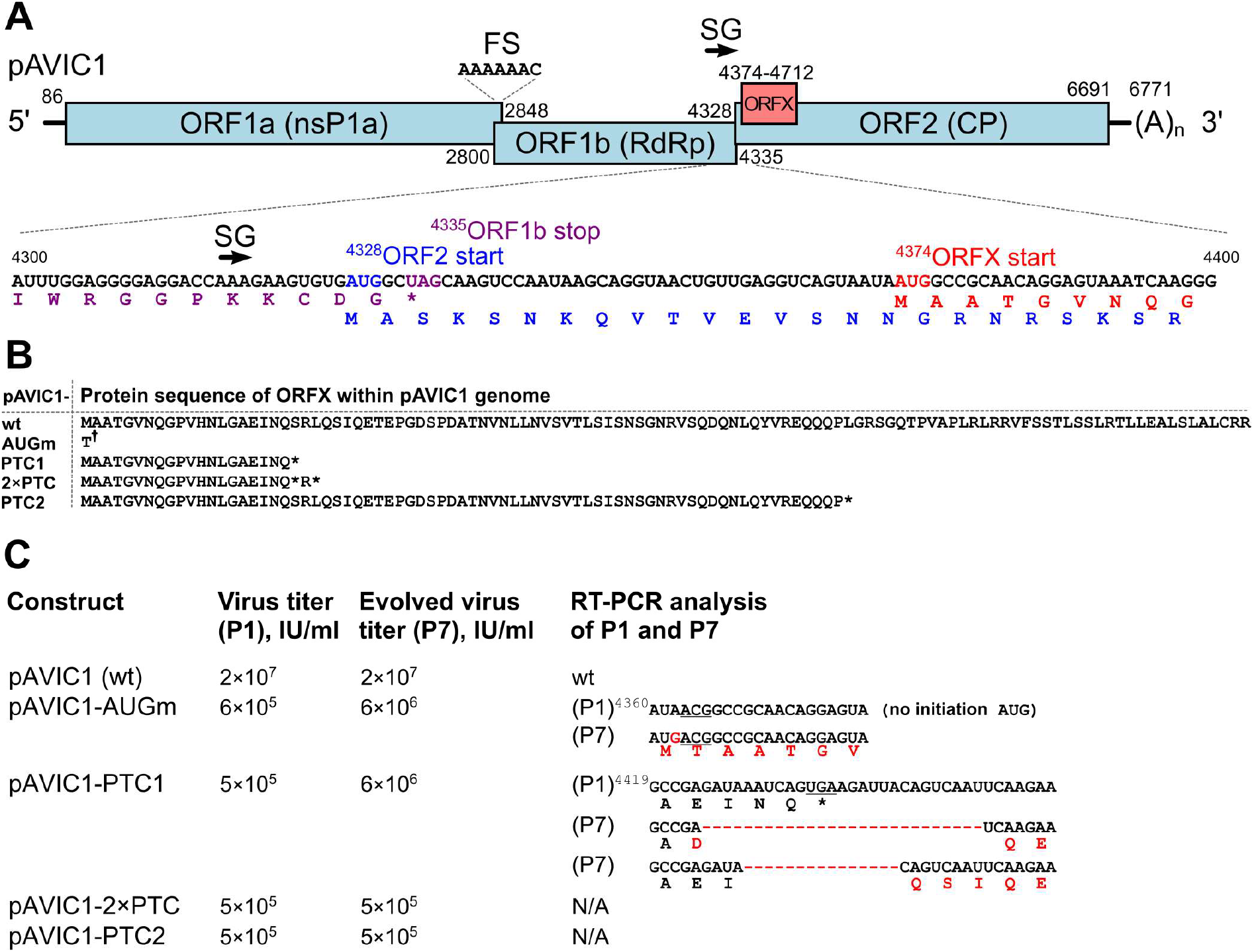
Design and properties of XP knockout viruses. **(A)** Schematic representation of the astrovirus genome. Numbers correspond to astrovirus genome nucleotides in the pAVIC1 infectious clone; FS, frameshift signal; SG, subgenomic promoter. The sequence of nucleotides 4300–4400 is shown below, indicating the positions of overlapping elements and the corresponding translated proteins: ORF1b (purple), ORF2 (blue) and ORFX (red). **(B)** XP amino acid sequences for the wt and mutant pAVIC1-derived viruses († see main text for note on the AUGm mutant). **(C)** Titers (infectious units per ml, IU/ml) of recombinant viruses after RNA transfection followed by first passage (P1) and after seven passages (P7) in Caco2 cells. RT-PCR analysis of viral RNA isolated after P1 and P7 for selected recombinant viruses.

### XP is essential for virus release but has only a minor effect on RNA replication

ORFX overlaps ORF2, which encodes the structural polyprotein, and therefore the expression of XP is likely required in late stages of the virus replication cycle. To rule out the possibility of XP directly influencing viral RNA replication, we developed a replicon system comprising an intact astrovirus genome up to the end of ORFX, followed by a 2A-RLuc cassette fused in either the ORF2 or ORFX reading frame, followed by the last 624 nt of the virus genome and a poly-A tail (Fig. 4A). These replicons provide a direct measurement of translated product associated with activity of the subgenomic promoter. Two ORFX knock-out mutations were copied in both versions of the replicon to evaluate the significance of XP for RNA replication (Fig. 4B). Experiments were performed in BSR and Huh7.5.1 cell lines, both of which have been reported to support astrovirus replication ^16,17^. Replication was confirmed by an 1200– 2100 fold difference in relative luciferase activity between wt and an RdRp knockout mutant (GDD → GNN) at 9 and 12 hours post transfection (Fig. 4C, Fig. S12A). The mutations introduced to knock out ORFX had minor effect on luciferase activity when 2A-RLuc was fused in the ORF2 reading frame (Fig. 4C, Fig. S12A). Consistent with our Ribo-Seq results (Fig. 2C), luciferase activity for the ORFX reading frame was 14% (BSR cells; Fig. 4C) and 21% (Huh7.5.1 cells; Fig. S12A) of luciferase activity for the ORF2 reading frame. Thus, ORFX is efficiently translated but has only a minor (if any) direct effect on viral RNA replication.

**Figure 4.**
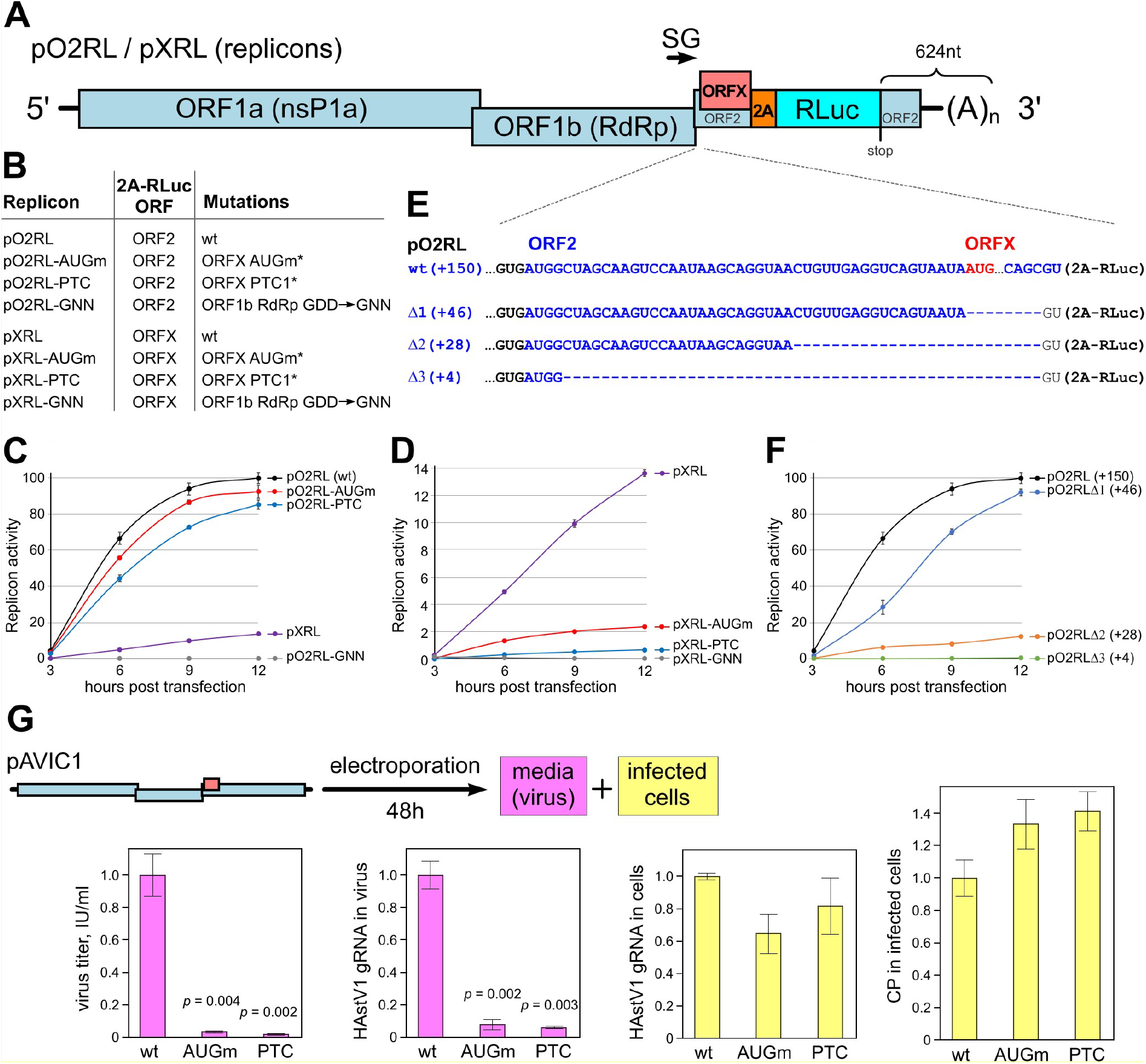
Analysis of replication stages affected by XP. **(A)** Schematic representation of the astrovirus replicon. The 2A-RLuc cassette is fused in either the ORF2 (pO2RL) or ORFX (pXRL) reading frame. **(B)** List of replicon mutants showing the RLuc reading frame and introduced mutations. * See Fig. S11 for exact sequences. **(C, D)** Relative replicon luciferase activities measured after RNA transfection of BSR cells (see Fig. S12 for Huh7.5.1 cells). Values are normalized so that the maximum wt value is 100%. **(E)** Schematic representation of deletion mutants in the pO2RL replicon; ORF2 (blue), ORFX initiation codon (red). **(F)** Relative replicon luciferase activities of deletion mutants. **(G)** Schematic design and results of experiment to quantify virus titer, RNA and protein levels in released virions and infected cells. BSR cells were electroporated with pAVIC1-wt, -AUGm or -PTC T7 RNAs and incubated for 48 h until appearance of CPE. Clarified supernatants were titrated (first graph) or treated with RNase I and used for viral RNA isolation and subsequent quantification by qRT-PCR (second graph). Total RNA from infected cells was isolated and virus gRNA was quantified by qRT-PCR (third graph). Aliquots of electroporated cells were seeded on a 96-well plate, incubated for 48 h, fixed, permeabilized, stained with anti-CP antibody, and imaged by LI-COR followed by quantification using LICOR software (fourth graph). Graphs show means ± s.d. from n = 3 biologically independent experiments (panels C, D, F, G).

As expected, both XP knockout mutations resulted in a substantial drop in ORFX-frame luciferase activity (Fig. 4D, Fig. S12B). This drop was more pronounced for the PTC mutant (5.5±0.7% wt in BSR cells and 4.1±1.7% in Huh7.5.1 cells) than for the AUGm mutant (AUG to ACG; 17.9±0.1% wt in BSR cells and 13.2±0.3% in Huh7.5.1 cells). ACG codons are known to permit initiation when in a strong initiation context (e.g. A at −3 and G at +4 as in the AUGm mutant) ^18^. Since we wished to only use mutations that were synonymous in ORF2, we were unable to mutate the ORFX AUG codon in any other way. When utilized as an initiation codon, ACG is expected to be decoded as methionine by initiator Met-tRNA. Therefore the AUGm mutant is still expected to produce wt XP, albeit at a greatly reduced level.

The astrovirus subgenomic promoter is also situated within this region of the astrovirus genome (Fig. 4A). Although effective replication occurred even in the absence of XP translation (Fig. 4C), the introduced mutations could still have a potential effect on sgRNA production via alteration of RNA structure and/or other interactions. To map the minimal region required for wt levels of sgRNA production in the context of the pO2RL replicon, we gradually truncated the sequence between the start of ORF2 and ORFX. Surprisingly, we found that a replicon containing only the first 4 or 28 nucleotides of ORF2 has very poor subgenomic reporter activity (0.6±0.03 and 12.5±0.1% that of wt pO2RL, respectively). In contrast, including the first 46 nucleotides of ORF2 restored subgenomic reporter activity to 92% of wt (Fig. 4F, S12C). These data suggest that the astrovirus subgenomic promoter is significantly longer than previously reported ^19^ and extends into the 5′ part ORF2, but does not appear to extend into ORFX.

To determine which stage of the virus replication cycle is affected in XP knock-out viruses, we quantified released virus and viral protein and RNA levels for BSR cells electroporated with *in vitro* transcribed full-length pAVIC1 T7 RNAs. As expected, neither RNA nor protein levels were affected in cell-derived samples (Fig. 4G). In contrast, in media-derived samples, XP mutant titers were significantly below wt titers when analyzed by qRT-PCR (fold differences = 0.08 and 0.05, *p* = 0.002 and 0.003 for AUG vs wt and PTC vs wt respectively; 2-tailed *t*-tests with separate variances) or virus titration (Fig. 4G). These analyses indicate the involvement of XP late in the virus replication cycle, with XP potentially acting in either virus assembly or virus release. Since previous structural studies on the HAstV CP ^20^ and virion ^21^ have not indicated the presence of any additional proteins in the compact 43-nm virion structure, the more likely role for XP is in virus release.

### XP localizes to plasma membrane and perinuclear membranes with an extracellular N-terminus topology

To further investigate the function of XP, we studied its intracellular localization. To make the small XP protein more stable and enable visualization in transfected cells, we fused it either N- or C-terminally with mCherry in the context of a mammalian expression vector. The diffuse cytoplasmic localization of mCherry was drastically affected in both fusions, where it was relocalized to plasma and perinuclear membranes (Fig. 5A). This strongly suggests the presence of a membrane-interacting domain in XP, which was not predicted by TM domain prediction software (Fig. S8; see Methods). Since ORFX is completely embedded within ORF2, it may have greatly decreased evolutionary flexibility, perhaps resulting in the evolution of a non-canonical TM region. Some of the putative XP proteins encoded by other astroviruses do in fact have predicted TM regions (Fig. S9). We also confirmed membrane and nuclear association of the XP-fused mCherry proteins by subcellular fractionation of transfected HeLa cells and subsequent analysis of the fractions (Fig. 5B). To investigate the potential topology of XP within the plasma membrane, we probed live electroporated HeLa cells with anti-mCherry antibody, fixed the cells and analyzed them by confocal microscopy. We observed punctate staining across the plasma membrane when mCherry was fused to the N-terminus but not when it was fused to the C-terminus, suggesting an extracellular N-terminal topology and potential multimerization (Fig. 5C-D).

**Figure 5.**
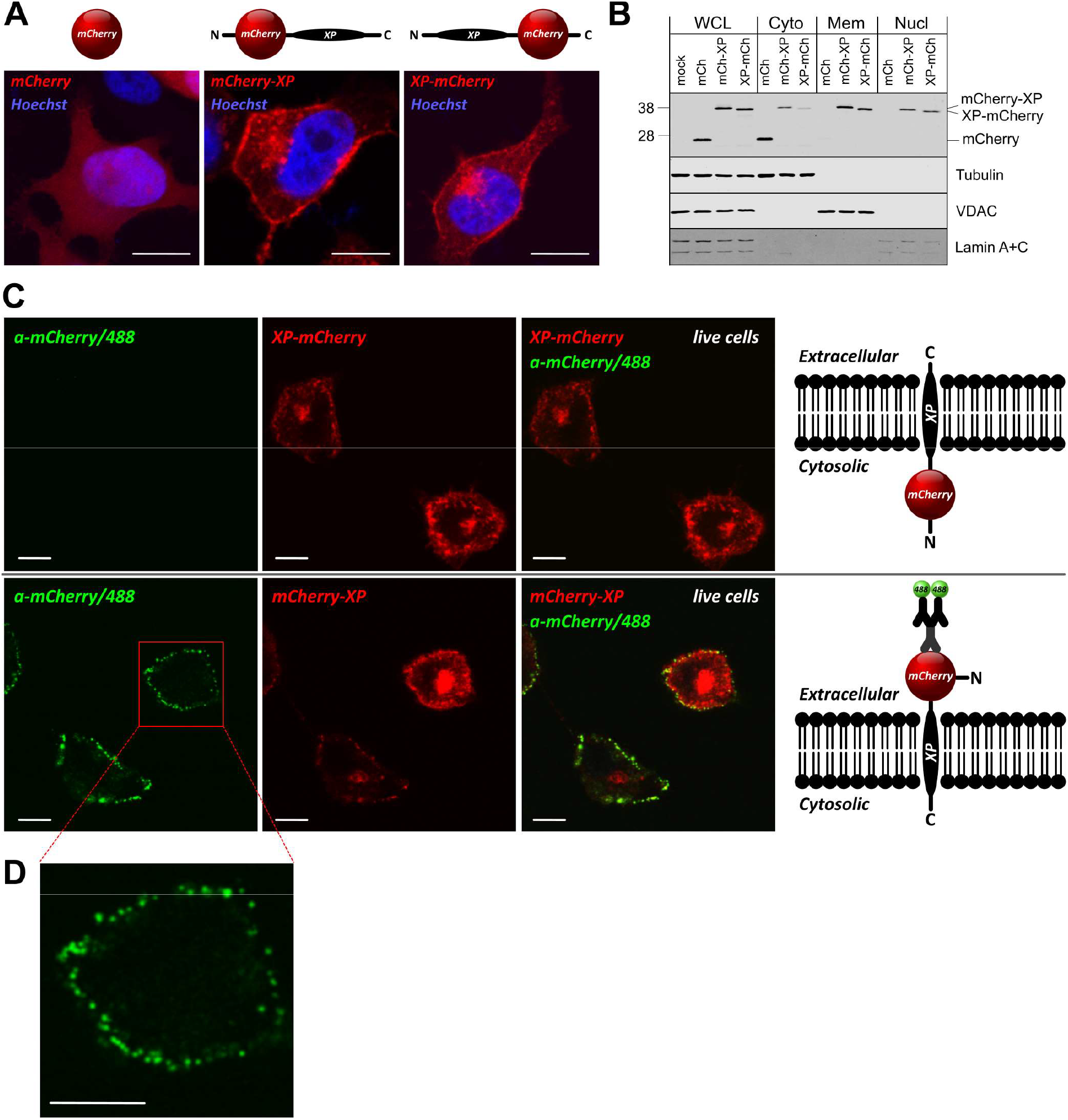
Cellular localization and membrane topology of XP. HeLa cells were electroporated with pCAG-mCherry, pCAG-mCherry-XP or pCAG-XP-mCherry. **(A)** Representative confocal images of fixed and permeabilized cells were visualized for mCherry (red) and stained for nuclei (Hoechst, blue). The images are averaged single plane scans. **(B)** Cell lysates were fractionated and whole cell lysate (WCL), cytoplasmic (Cyto), membrane (Mem) and soluble nuclear (Nucl) fractions were analyzed by immunoblotting with antibodies to mCherry, tubulin, VDAC or Lamin A+C as indicated. See Fig. S13 for complete images. **(C)** Live cells after electroporation were probed with anti-mCherry antibody, fixed, and visualized using confocal microscopy. The images are averaged single plane scans. **(D)** Zoom-in image of punctate structures seen in C. All scale bars are 10 µm (A, C, D).

### XPs from HAstV1 and related astroviruses have a viroporin-like activity

Given the above results, we hypothesized a possible viroporin function for XP. Viroporins are virus-encoded ion channel proteins which disturb membrane integrity leading to permeabilization. Given the C-terminal below-threshold TM predictions for HAstV XPs (Fig. S8), we hypothesized that the C-terminal region might harbour a TM pore-forming domain. Additional support came from Kyte-Doolittle hydrophobicity profiles (Fig. 6B) and a C-terminal α-helical structure prediction for HAstV1–8 XPs (Fig. S14). A helical wheel representation of this region revealed a penta-leucine hydrophobic face in HAstV1 (Fig. 6A). Although these leucines are not entirely conserved between the eight HAstV serotypes (Fig. 1F), each XP was still predicted to contain an amphipathic α-helix in this region ^22^. To test for viroporin-like activity, we utilized a previously described Sindbis virus replicon (SINV repC) ^23^ to overexpress XP or control proteins in mammalian cells. This replicon has been demonstrated to be an extremely useful tool for investigating viroporins from different RNA viruses via induced permeabilization of the plasma membrane in BHK cells to the translation inhibitor hygromycin B (HB). BSR cells (a clone of BHK cells) were electroporated with *in vitro* transcribed SINV repC RNAs, and new protein synthesis was labelled with L-azidohomoalanine (AHA) in the presence or absence of HB, followed by lysis and click chemistry-based on-gel detection ^24^. Expression of XP or Strep-tagged enterovirus 2B (a well-characterized viroporin ^25^) led to almost total inhibition of protein synthesis (Fig. 6C-D) indicating that both proteins induced pronounced cell permeabilization to HB.

**Figure 6.**
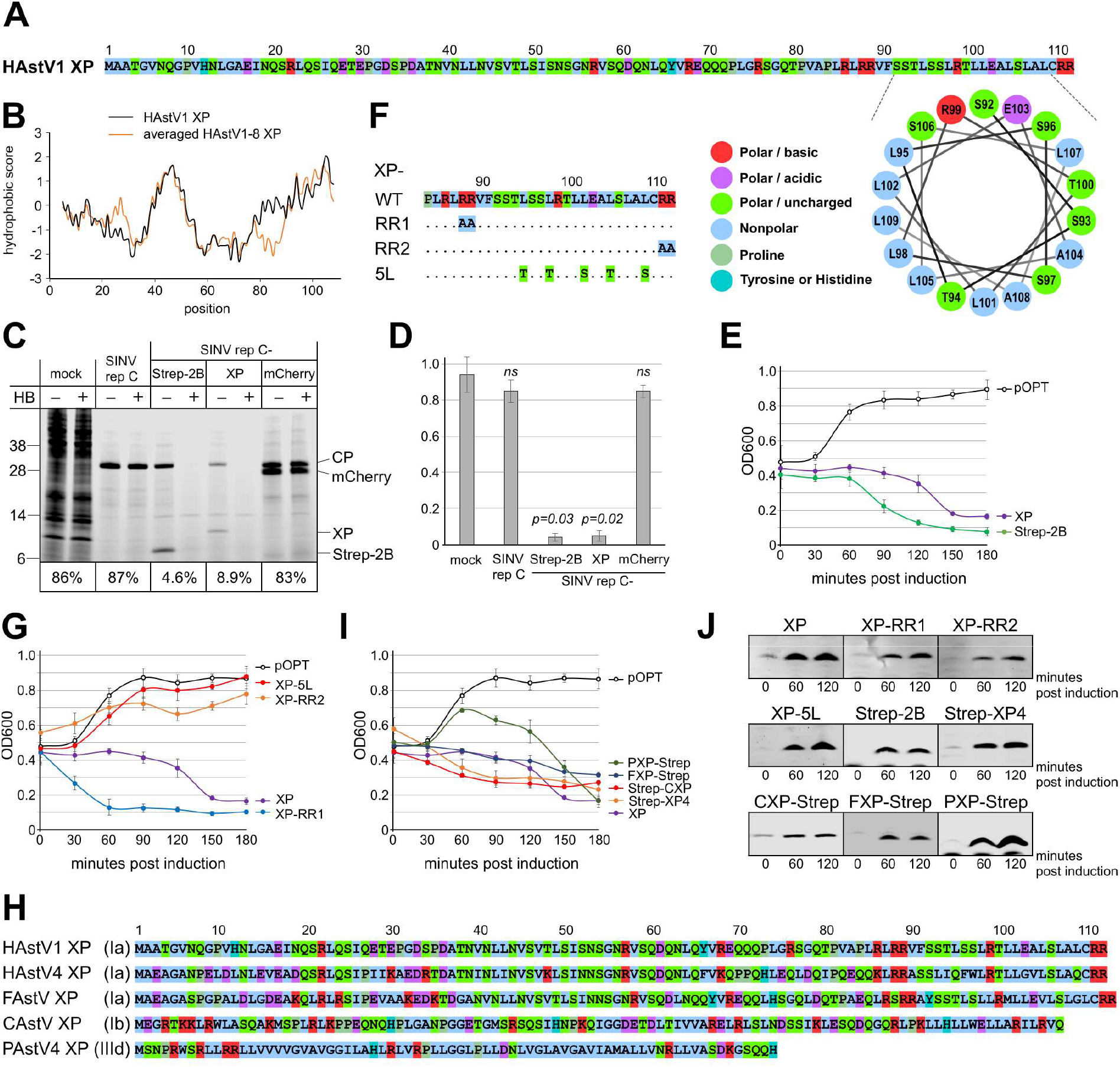
XP from HAstV1 and related astroviruses has viroporin-like activity. **(A)** HAstV1 XP sequence and helical wheel representation of amino acids 92–109. **(B)** Kyte-Doolittle hydropathy plots of HAstV XPs (see Fig. S16 for individual plots). **(C)** Membrane permeabilization in BSR cells at 8 h post RNA electroporation with Sindbis virus replicons (SINV repC) expressing HAstV1 XP, enterovirus Strep-2B (positive control) or mCherry (negative control). Ongoing protein synthesis was labelled with 1 mM AHA in the presence or absence of 1 mM HB as a translation inhibitor. Cells were lysed and AHA-bearing proteins were ligated to the fluorescent reporter IRDye800CW Alkyne by click chemistry, separated by SDS-PAGE, and visualized by in-gel fluorescence. The numbers below each pair of samples indicate protein synthesis quantified for HB-treated cells relative to the values obtained for untreated cells. **(D)** Statistical analysis of membrane permeabilization caused by XP and the indicated control proteins in BSR cells. Bars indicate the amount of protein synthesis in HB-treated cells relative to untreated cells (mean ± s.d.; *n* = 3 biologically independent experiments). (E,G,I) *E. coli* lysis assay for HAstV1 XP **(E)** and indicated mutants **(G)** or XPs derived from other astrovirus species **(I)** (mean ± s.d.; *n* = 3 biologically independent experiments). **(F)** Amino acid mutations used in G. **(H)** Astrovirus XP sequences tested in I. **(J)** Western blots for E, G and I confirming expression of the relevant products (see Fig. S17 for full images).

Another widely used assay to assess the ability of proteins to permeabilize cellular membranes is based on impaired growth of *Escherichia coli* upon induced overexpression of a membrane-permeabilizing protein ^26,27^. Consistent with the results observed in the mammalian system, induced expression of XP or the enteroviral viroporin 2B in *E. coli* resulted in cytotoxicity and impaired growth (Fig. 6E). There are two features that are generally associated with viroporin membrane permeabilizing activity: (i) an amphipathic α-helix that facilitates oligomerization to form the TM pore, and (ii) adjacent positively charged residues that anchor the viroporin in the membrane ^27,28^. To test for these features in XP, two conserved RR motifs present in all eight HAstV serotypes (Fig. 1F) were mutated to alanines (Fig. 6F). Interestingly, only the mutation of the very C-terminal arginines resulted in loss of XP activity in this system (Fig. 6G). Mutation of the hydrophobic leucine stretch (L95, L98, L102, L105 and L109, Fig. 6A) to serines and threonines (Fig. 6F) also resulted in complete loss of XP activity (Fig. 6G). Thus, the C-terminal domain of XP is involved in membrane permeabilization and harbours key features of a viroporin.

To investigate conservation of XP activity across different astrovirus species, we tested several other astrovirus XPs in the bacterial lysis assay. We used XPs from closely related HAstV4, feline (FAstV; genotype Ia), canine (CAstV; genotype Ib) and distantly related porcine (PAstV4; genotype IIIc) astroviruses in this experiment (Fig. 6H). Protein expression constructs were created to express XPs with N- or C-terminal Strep tags, and combinations resulting in detectable protein were chosen for the assay. Induction of each of the four XPs resulted in strong inhibition of bacterial growth (Fig. 6I), with XP expression confirmed by western blot (Fig. 6J). Similar to HAstV XPs, all tested proteins contain one or two predicted amphipathic α-helices (Fig. S15). Thus, these findings indicate that this newly identified astrovirus protein has a membrane-permeabilizing function.

## DISCUSSION

The data presented here demonstrate the existence of an additional protein, XP, encoded within the human astrovirus genome. XP is important for virus growth, localizes to the plasma membrane, and plays a role in virus release. Viroporins have been reported for many enveloped and non-enveloped viruses and – although they can play roles in virus entry and modulation of cellular pathways – most often they facilitate virus assembly or release ^26^. However, no viroporin candidate had been previously predicted for astroviruses. Identifying a viroporin is challenging due to the lack of homology among viroporins from different viruses. Viroporins are typically small hydrophobic integral membrane proteins of around 100 aa in size, with two motifs – an amphipathic α-helix and an adjacent cluster of positively charged residues; mutation of these residues generally abolishes viroporin activity ^27^. Using two different assays and computational approaches, we show that XP fulfils these criteria. XP is capable of permeabilizing cellular membranes and has a distinct N-terminal extracellular topology with one TM domain, thus making XP a candidate class IA viroporin, a class which also includes the influenza A virus M2, coronavirus E and HIV-1 Vpu proteins ^26,28^. Future work will be needed to confirm potential ion channel activity, and characterize ion specificity, structural organisation, and processes affected by XP expression in the context of viral infection. The localization of XP not only at the plasma membrane but also in the perinuclear region raises the possibility that XP may also have additional functions. Whereas the C-terminal α-helix appears to be associated with the cell-permeabilizing activity, additional functions (if any) of the extended N-terminal domain of XP remain to be studied.

Comparative genomic analysis suggests that the presence of a protein-coding ORF overlapping ORF2 is widespread in mammalian astroviruses. It appears to be almost ubiquitous in genogroups I, III and IV and is probably present in the single genogroup V sequence. It also appears to be present in various unassigned sequences and clades. ORFX frequently has a hydrophobic stretch that is often predicted to be a TM domain and in other cases may represent a non-canonical TM domain refractory to detection with standard TM-predicting software (as appears to be the case for HAstV1). While normally absent from genogroup II astroviruses, an ORFX appears to be present in one clade (herein referred to as IId), whereas in the clade referred to herein as IIc we predicted an ORFY in the −1 instead of +1 frame, and accessed via ribosomal frameshifting instead of leaky scanning. Given the sporadic appearance of an overlapping ORF across the genogroup II phylogeny, it seems likely that the genogroup II ORFX and ORFY evolved independently from ORFX in genogroups I, III and IV, and may also therefore have different functions. In contrast, these other ORFXs may or may not have a common ancestor and/or common function (it is unclear to what extent genogroups I, III and IV form a monophyletic group; Fig. S1, Fig. S2). The N-terminal ~70 aa of CP are dispensable for particle assembly and likely structurally disordered ^20^ and may therefore be evolutionarily fairly flexible. Thus, this region of ORF2 may be unusually tolerant to the coding constraints imposed by overlapping genes. Together with the high translation level of sgRNAs at later timepoints (Fig. 2A), and the ease with which 5′ proximal ORFs can be expressed (requiring only leaky scanning rather than more complex expression mechanisms such as internal ribosome entry or ribosomal frameshifting ^10^), the 5′ region of the sgRNA may be particularly well-suited to the evolution of an overlapping gene, consistent with multiple independent origins of ORFXs.

During this study we also developed and characterized the first astrovirus replicon system. This will be of broad utility to the astrovirus research community. Astroviruses are one of the major causes of infant gastroenteritis; they are widespread among mammals; and non-classical human-infecting astroviruses (such as the MLB and VA/HMO clades) have recently been recognized. Nonetheless, despite their ubiquity and importance, astroviruses represent some of the least well-studied human viruses, partly because it has been difficult to establish efficient lab systems to study them. The replicon system developed herein will now permit detailed characterization of astrovirus replication and gene expression, and facilitate research into antiviral drugs.

In summary, using comparative genomic analyses we predicted a new gene in the *Mamastrovirus* genus; using ribosome profiling we demonstrated XP expression in HAstV1-infected cells; and using a range of techniques we demonstrated the crucial role of XP in virus growth by promoting virus release, which is likely associated with its membrane-permeabilizing activity. These findings add a new dimension to astrovirus molecular biology, with potential impacts for new therapeutics (e.g. compounds that block XP activity) and vaccine development (e.g. by inhibiting XP expression).

## MATERIALS AND METHODS

### Comparative genomic analysis

Mammalian astrovirus nucleotide sequences were downloaded from the National Center for Biotechnology Information (NCBI) on 26 July 2018. Patent sequence records and sequences with ≥20 ambiguous nucleotide codes (e.g. “N”s) were removed. For the full-genome analyses, only sequences covering all or nearly all of ORF1a, ORF1b and ORF2 were retained, giving 221 sequences (listed in Fig. S1). For the ORF2 analyses, only sequences covering all or nearly all of ORF2 were retained, giving 415 sequences (listed in Supplementary Dataset 1). To identify the correct 5′ end of ORF1b, we identified the AAAAAAC frameshift site. To identify the correct initiation site of ORF2, we identified the highly conserved sgRNA promoter nucleotides ^29^ and selected the next ORF2-frame AUG codon as the ORF2 start site in representative reference sequences; for the other sequences, the ORF2 start site was identified by amino acid alignment to one of the reference sequences. ORF1b and ORF2 sequences were extracted, translated to amino acid sequences, aligned with MUSCLE ^30^, and maximum likelihood phylogenetic trees were estimated using the Bayesian Markov chain Monte Carlo method implemented in MrBayes version 3.2.3 ^31^ sampling across the default set of fixed amino acid rate matrices, with 1,000,000 (221-sequence trees; Fig. S1 and Fig. S2) or 5,000,000 (415-sequence tree; Fig. S4) generations, discarding the first 25% as burn-in (other parameters were left at defaults). Trees were visualized with FigTree (http://tree.bio.ed.ac.uk/software/figtree/).

The 221-sequence ORF1b tree was used to manually select clades (Fig. 1B) for full-genome SYNPLOT2 analyses (Fig. S3). For the ORF2-only SYNPLOT2 analyses (Fig. S5), we used a more objective method to select clades. Through an iterative procedure of clustering the 415 ORF2 sequences based on amino acid identity to a set of reference sequences, and selecting sequences distal from all reference sequences as new reference sequences, we arrived at a set of 42 ORF2 reference sequences: KT946734, Z25771, JN420356, KP404149, JX556691, AY179509, Y15937, FJ973620, JF713713, JX544743, JF713710, LC047798, FJ222451, JX556693, KT946736, KP663426, KY855439, JF729316, KY855437, KY024237, MG693176, JF755422, EU847155, KT946731, LC047794, KF787112, JN420352, JN420359, LC047787, KT963069, HQ916316, KX645667, KT963070, FJ890355, HQ668129, HQ668143, FJ571066, EU847145, EU847144, FJ571067, GQ415660, FJ571068. The highest reference:reference pairwise CP amino acid identity is 0.5173; no non-reference has <50% identity to a reference, and only 20 have <55% identity; 82 and 4 sequences have >50% and >55% identity, respectively, to >1 references. Each non-reference sequence was then clustered with the reference sequence to which it has highest CP amino acid identity. This resulted in 16 singleton clusters and 26 multi-sequence clusters.

Synonymous site conservation was analyzed with SYNPLOT2 ^9^. For the full-genome analyses we generated codon-respecting alignments using a procedure described previously ^9^. In brief, each individual genome sequence was aligned to a reference sequence using code2aln version 1.2 ^32^. Genomes were then mapped to reference sequence coordinates by removing alignment positions that contained a gap character in the reference sequence, and these pairwise alignments were combined to give the multiple sequence alignment. To assess conservation at synonymous sites, the ORF1a, ORF1b and ORF2 coding regions were extracted from the alignment (with codons selected from the longer ORF in each overlap region), concatenated in-frame, and the alignment analyzed with SYNPLOT2 using a 25-codon sliding window. Conservation statistics were then mapped back to reference genome coordinates for plotting. For the ORF2-only SYNPLOT2 analyses, any duplicate sequences were removed and the remaining ORF2 sequences in each clade were translated, aligned using MUSCLE as amino acid sequences, back-translated to codon-respecting nucleotide alignments, and the alignment analyzed with SYNPLOT2 as above. In contrast to the full-genome alignments, all alignment gaps were retained instead of mapping to a specific reference sequence coordinate system. Calculation of the pI and molecular mass of XP peptides and other sequence processing were performed with pepstats and other programs from the EMBOSS package ^33^. TM domains were predicted with Phobius (EMBL-EBI) ^34^. The XP proteins of HAstVs 1–8 were additionally queried with TMHMM (http://www.cbs.dtu.dk/services/TMHMM/; weak TM prediction for HAstV3, no TM predicted for other HAstVs) and SOSUI (http://harrier.nagahama-i-bio.ac.jp/sosui/sosui_submit.html; no TMs predicted). XP secondary structures were predicted with RaptorX ^35^. To search for potential homologues of XP, XPs from Fig. S6D, Fig. S7 and Fig. S9 were queried with HHpred ^33^. Only one of the 33 queries retrieved a match with e-value < 1 – namely murine astrovirus (genogroup IIIa) which obtained an e-value 0.38 hit to a fragment of Protein Data Bank accession 5NIK_K, an *E. coli* outer membrane ion channel protein (most likely a manifestation of convergent evolution).

### Cells and viruses

BSR (single cell clone of BHK-21 cells) and HeLa cells (ATCC, CCL-2) were maintained at 37 °C in DMEM supplemented with 10% fetal bovine serum (FBS), 1 mM L-glutamine, and antibiotics. Huh7.5.1 cells ^34^ (Apath, Brooklyn, NY) were maintained in the same media supplemented with non-essential amino acids. All cells were mycoplasma tested (MycoAlert^TM^ PLUS Assay, Lonza); BSR and Caco2 cells were also tested by deep sequencing.

The infectious clone of HAstV1 (pAVIC1, GenBank accession number L23513.1) was described previously ^16^. The reverse genetics procedure was compiled from several previously published approaches ^16,17^. Initial virus was recovered from T7 transcribed RNA using reverse transfection of Huh7.5.1 cells by Lipofectamine® 2000 (Invitrogen) (for virus rescue, see Fig. 3) or electroporation of BSR cells in PBS at 800 V and 25 µF using a Bio-Rad Gene Pulser Xcell^TM^ electroporation system (see Fig. 4 for an analysis of virus RNA in cells versus released particles). For virus passaging, the collected supernatant was treated with 10 µg/ml trypsin (Type IX, Sigma, #T0303) for 30 min at 37 °C, diluted 5 times with serum-free media, and used for infection of Caco2 cells. After 3 h of incubation, the virus containing media was replaced with serum free media containing 0.6 µg/ml trypsin, and cells were incubated for 48–72 h until appearance of CPE. After 3 freeze-thaw cycles, the viral stocks were aliquoted, frozen and stored at −70 °C. Viral stocks were titrated as previously described ^35^, but using infrared detection readout, combined with automated LI-COR software-based quantification.

### Virus evolution

Virus evolution was performed in Caco2 cells in triplicate by passaging mutant viruses at different dilutions (corresponding to MOIs ranging from 0.001 to 0.1). Since HAstVs do not form plaques in Caco2 cells, an end-point dilution approach was used instead of plaque purification. Infections that resulted in CPE by 5 dpi were used for subsequent passage at a similar dilution and incubation time. Following an increase of at least 10-fold over P1 titer (Fig. 3C), individual flasks with infected Caco2 cells were used for RNA isolation by Direct-zol RNA MicroPrep (Zymo research), RT-PCR and Sanger sequencing of the fragment containing the mutated region of the virus genome.

### Plasmids

For mammalian expression of XP, the coding sequence of mCherry alone or HAstV1 XP fused to mCherry at either the N- or C-terminus was inserted into vector pCAG-PM ^36^ using *Afl*II and *Pac*I restriction sites. The resulting constructs – designated pCAG-mCherry, pCAG-XP-mCherry, and pCAG-mCherry-XP – were confirmed by sequencing.

All virus genome mutations (Fig. S11) were introduced using site-directed mutagenesis of pAVIC1 and confirmed by sequencing. The resulting plasmids were linearized with *Xho*I prior to T7 RNA transcription. To create the HAstV1 replicon system, the pAVIC1 infectious clone was left intact up to the end of ORFX, then followed by a foot and mouth disease virus 2A sequence and a *Renilla* luciferase (RLuc) sequence fused in either the ORF2 (pO2RL) or ORFX (pXRL) reading frame, followed by the last 624 nt of the virus genome and a poly-A tail (Fig. 4A). All mutations were introduced from the corresponding pAVIC1 mutants using available restriction sites and all constructs were confirmed by sequencing. The resulting plasmids were linearized with *Xho*I prior to T7 RNA transcription.

For bacterial expression of XP and related control proteins (mCherry and N-terminally Strep-tagged 2B from Echovirus 7), the relevant coding sequences were inserted into the pOPT expression plasmid ^37^ between NdeI and *Bam*HI restriction sites. Feline (KF374704.1), porcine (LC201600.1) and canine (FM213332.1) XP coding sequences were commercially synthesized (Integrated DNA Technologies). Sequence encoding HAstV4 XP was RT-PCR amplified from a clinical HAstV4 isolate kindly provided by Susana Guix (University of Barcelona, Spain). Each XP coding sequence was inserted into the pOPT plasmid with a C- or N-terminal Strep-tag as indicated (Fig. 6I). For the permeabilization assay, Sindbis virus derived replicons (SINV repC) expressing XP, mCherry or Strep-tagged enterovirus (echovirus 7) 2B were created exactly as previously described ^23^.

### Ribosome profiling

Caco2 cells were grown on 150-mm dishes to reach 80–90% confluency. The cells were infected at MOI 5 with HAstV1 virus stock (passage 2, derived from pAVIC1 T7 RNA). At 12 hpi, cells were either not treated (NT) or treated with 50 µM LTM, flash frozen in a dry ice/ethanol bath, and lysed in the presence of 0.36 mM cycloheximide. Cell lysates were subjected to Ribo-Seq based on the previously described protocols ^14,38^, except Ribo-Zero Gold rRNA removal kit (Illumina), not DSN, was used to deplete ribosomal RNA. Amplicon libraries were deep sequenced using an Illumina NextSeq platform.

### Computational analysis of Ribo-Seq data

Ribo-Seq analysis was performed as described previously ^14^. Adaptor sequences were trimmed using the FASTX-Toolkit (http://hannonlab.cshl.edu/fastx_toolkit) and trimmed reads shorter than 25 nt were discarded. Reads were mapped to host (*Homo sapiens*) and virus RNA using bowtie version 1 ^39^, with parameters -v 2 --best (i.e. maximum 2 mismatches, report best match). Mapping was performed in the following order: host rRNA, virus RNA, host RefSeq mRNA, host non-coding RNA, host genome.

To normalize for library size, reads per million mapped reads (RPM) values were calculated using the sum of positive-sense virus and host RefSeq mRNA reads as the denominator. A +12 nt offset was applied to the RPF 5′ end positions to give the approximate ribosomal P-site positions. To calculate the phasing and length distributions of host and virus RPFs, only RPFs whose 5′ end (+12 nt offset) mapped between the 13th nucleotide from the beginning and the 18th nucleotide from the end of coding sequences (ORF1a, ORF1b and ORF2 for HAstV; RefSeq mRNAs for host) were counted, thus avoiding RPFs near initiation and termination sites. For Fig. S10, the dual-coding region where ORFX overlaps ORF2 was also excluded. Histograms of host RPF positions (5′ end +12 nt offset) relative to initiation and termination sites were derived from RPFs mapping to RefSeq mRNAs with annotated coding regions ≥450 nt in length and with annotated 5′ and 3′ UTRs ≥60 nt in length. Virus ORF1a, ORF1b and ORF2 ribosome densities (used herein as a proxy for translation levels) were calculated by counting RPFs whose 5′ end (+12 nt offset) mapped within the regions 101–2782, 2866–4311 or 4730–6676, respectively (i.e. excluding the dual-coding regions and excluding reads with P-sites mapping within 15 nt of initiation, termination or ribosomal frameshifting sites).

To compare phasing in the region of ORF2 overlapped by ORFX with the region of ORF2 not overlapped by ORFX (Fig. 2C) we counted RPFs whose 5′ end (+12 nt offset) mapped within the regions 4388–4693 or 4727–6673, respectively. We then compared the fraction of RPFs in phases 1, 2 and 3 in the ORF2/ORFX overlap region [*v*_1_, *v*_2_, *v*_3_] with the corresponding fractions for host mRNAs [*m*_1_, *m*_2_, *m*_3_], leading to the three equations [*v*_1_, *v*_2_, *v*_3_] ≈ *c* × [*m*_1_, *m*_2_, *m*_3_] + *x* × [*m*_3_, *m*_1_, *m*_2_], where *c* and *x* are the relative expression levels of ORF2 and ORFX, respectively, and *c* + *x* = 1. Writing [*q*_1_, *q*_2_, *q*_3_] = *c* × [*m*_1_, *m*_2_, *m*_3_] + (1 − c) × [*m*_3_, *m*_1_, *m*_2_] − [*v*_1_, *v*_2_, *v*_3_], we found *c* to minimize (*q*_1_^2^ + *q*_2_^2^ + *q*_3_^2^)^1/2^, giving *c* = 0.79 and 0.78 for repeats 1 and 2 respectively. Thus the estimated ORF2:ORFX expression ratios are 0.79:0.21 and 0.78:0.22, i.e. ORFX is expressed at ~27% of the level of ORF2.

### Permeabilization assay

BSR cells electroporated with RNA synthesized *in vitro* from the different linearized replicon constructs or with transcription buffer alone were seeded in 6-well plates. At 8 h post electroporation, cells were pretreated with 1 mM HB (Invitrogen) in methionine-free media (Gibco, Life Technologies) for 20 min or left untreated. Next, ongoing protein synthesis was labelled with 1 mM AHA (Invitrogen) in methionine free media in the presence or absence of 1 mM HB for 40 min. Finally, cells were lysed in 1% SDS containing PBS supplemented with protein inhibitor cocktail (Roche) and 250 U/ml bensonase (Sigma). Labelled proteins were ligated with 25 µM IRDye800CW Alkyne (LI-COR) in the presence of CuSO_4_ (CCT, 100 µM), tris-hydroxypropyltriazolylmethylamine (THPTA, CCT, 500 µM), aminoguanidine (Cambridge Bioscience, 5 mM) and sodium L-ascorbate (Sigma, 2.5 mM) for 2 h at room temperature. The final products were lysed, resolved in Novex 10–20% tricine protein gels (Invitrogen), fixed in 50% ethanol and 10% acetic acid for 10 min, scanned on LI-COR and analyzed by densitometry using LI-COR software. Experiments were repeated 3 times.

### *E. coli* lysis assay

For the assessment of protein viroporin-like activity in *E. coli, E. coli* BL21 (DE3) cells were transformed with pOPT constructs, grown in the presence of ampicillin (100 µg/ml) at 37 °C to an optical density at 600 nm (OD_600_) of 0.4 to 0.6, and then 1 mM isopropyl-b-D thiogalactopyranoside (IPTG) was added to induce protein expression. Subsequently, optical densities were measured for induced and non-induced samples in triplicate over a time course of 180 min post induction. Non-induced, and 60 and 120 min post induction samples were also collected for protein detection by western blot.

### Fractionation analysis

To analyze the cellular distribution of overexpressed XP fusions, electroporation of HeLa cells was performed in full media at 240 V and 975 µF using a Bio-Rad Gene Pulser. At 20 h post electroporation (hpe), cells were washed with PBS and fractionated using a subcellular protein fractionation kit for cultured cells (Thermo Scientific) according to the manufacturer’s instructions. Equal aliquots of whole cell lysate, cytoplasmic, membrane and soluble nuclear fractions were analyzed by western blot using the indicated virus- or cellular target-specific antibodies.

### Immunofluorescence microscopy

For the analysis of intracellular localization of overexpressed XP fusions, electroporation of HeLa cells was performed as described in the previous section. At 16 hpe, cells were fixed with 4% paraformaldehyde (PFA) for 20 min at room temperature, followed by permeabilization with PBS containing 0.1% Triton X-100 for 10 min. Nuclei were counter-stained with Hoechst (Thermo Scientific). For life cell probing, at 16 hpe cells were washed with cold PBS and incubated with anti-mCherry antibody (abcam, ab167453, 1:500 dilution) on ice for 1 h, then washed with PBS, fixed with 4% PFA for 20 min at room temperature and incubated with secondary antibody (Alexa Fluor 488-conjugated goat anti-rabbit, Thermo Fisher, A21441). The images are single plane images taken with a Leica SP5 Confocal Microscope using a water-immersion 63× objective.

### Analysis of cellular and released virus samples

BSR cells were electroporated with *in vitro* transcribed RNAs derived from wt or mutant pAVIC1 using a double-pulse protocol in a BioRad Gene Pulser at 800 V and 25 µF in PBS. After 4 hpe, cells were washed 3 times and supplemented with virus production serum-free media (VP-SFM, Gibco) containing 0.6 µg/ml trypsin. At 48 hpe media samples were collected and centrifuged at 9,600 g for 5 min. The released virus samples were titrated on Caco2 cells as described above. For RT-qPCR analysis, a 150 µl aliquot of each sample was mixed with 4 × 10^6^ plaque forming units (PFUs) of purified Sindbis virus (SINV) stock, which was used for normalization and to control the quality of RNA isolation. The virus samples were treated with 300 units of RNase I (Ambion) for 30 min followed by SUPERaseI RNase Inhibitor (Thermo Fisher) to eliminate RNA contaminants outside of virus particles. RNA was then extracted using the Qiagen QIAamp viral RNA mini kit. Reverse transcription was performed using the QuantiTect reverse transcription kit (Qiagen) using virus-specific reverse primers for SINV (GTTGAAGAATCCGCATTGCATGG) and HAstV1 (TACTGCTGTAGCAATAAGGCCACG). HAstV1 (TGCTATTGGTACTGTCATGGG and GGTGTGAAATGGAATTGTGGG) and SINV (GAAACAATAGGAGTGATAGGCA and TGCATACCCCTCAGTCTTAGC) specific primers were used to quantify corresponding virus RNAs; the primer efficiency was within 95–105%. Quantitative PCR was performed in triplicate using SsoFast EvaGreen Supermix (Bio-Rad) in a ViiA 7 Real-time PCR system (Applied Biosystems) for 40 cycles with two steps per cycle. Results were normalized to the amount of SINV RNA in the same sample. Fold differences in RNA concentration were calculated using the 2^−ΔΔ*CT*^ method.

For HAstV1 CP quantification, the electroporated cells were seeded on 96-well plates at different dilutions, and at 24 hpe fixed with 4% PFA and permeabilized with 0.5% triton X-100 in PBS. CP was detected using Astrovirus 8E7 antibody (Santa Cruz Biotechnology) followed by LI-COR visualization and quantification.

For analysis of virus RNA present in infected cells, at 48 hpe cells were washed 3 times with PBS to eliminate extracellular virus and total RNA was extracted using Direct-zol RNA MiniPrep Plus (Zymo Research) according to the manufacturer’s protocol. Reverse transcription was performed using the QuantiTect reverse transcription kit (Qiagen) using random and HAstV1-specific reverse primers. For qPCR, hamster GAPDH-specific (GGCAAGTTCAAAGGCACAGTC and CACCAGCATCACCCCATTT) and HAstV1-specific primers (above) were used. Results were normalized to the amount of GAPDH RNA in the same sample. Fold differences in RNA concentration were calculated using the 2^−ΔΔ*CT*^ method.

### SDS-PAGE and immunoblotting

Lysates from the above mentioned assays were analyzed by SDS-PAGE, using standard 12% SDS-PAGE to resolve mCherry and its XP-fusion variants, and precast Novex™ 10–20% tricine protein gels (Thermo Fisher) to resolve XPs and enterovirus 2B. Proteins were then transferred to 0.2 µm nitrocellulose membranes and blocked with 4% Marvel milk powder in phosphate-buffered saline (PBS). Immunoblotting of mCherry was performed using anti-mCherry antibody (Abcam, ab167453). A custom rabbit polyclonal antibody raised against XP peptide SNSGNRVSQDQNLQ (GenScript; only able to detect strongly overexpressed XP) and an anti-Strep mouse antibody (Abcam, ab184224) were used for detecting HAstV1 XP and Strep-tagged proteins, respectively. The following antibodies were used for cellular targets: anti-tubulin (Abcam, ab15568), anti-VDAC1 (Abcam, ab14734), and anti-LAMIN A+C (Abcam, ab133256). Immunoblots were imaged and analyzed on a LI-COR imager. The original LI-COR scans are shown in Fig. S13 and Fig. S17.

### Reporter assay for astrovirus replicon activity

BSR and Huh7.5.1 cells were transfected in triplicate with Lipofectamine 2000 reagent (Invitrogen), using the protocol in which suspended cells are added directly to the RNA complexes in 96-well plates. For each transfection, 100 ng replicon plus 10 ng firefly luciferase-encoding purified T7 RNA (RNA Clean and Concentrator, Zymo research) plus 0.3 µL Lipofectamine 2000 in 20 µL Opti-Mem (Gibco) supplemented with RNaseOUT (Invitrogen; diluted 1:1,000 in Opti-Mem) were added to each well containing 10^5a^ cells. Transfected cells in DMEM supplemented with 5% FBS were incubated at 37 °C for the indicated times (Fig. 4C, D, F). Firefly and Renilla luciferase activities were determined using the Dual Luciferase Stop & Glo Reporter Assay System (Promega). Replicon activity was calculated as the ratio of Renilla (subgenomic reporter) to Firefly (cap-dependent translation, loading control), normalized by the same ratio for the wt O2RL replicon sequence. Three independent experiments each in triplicate were performed to confirm reproducibility of the results.

## ACKNOWLEDGEMENTS

We thank the Cambridge NIHR BRC Cell Phenotyping Hub for assistance with confocal microscopy. We thank Susanne Bell for technical assistance, Vanesa Madan, Alfredo Castello, Jia Lu, Ian Goodfellow and Myra Hosmillo for valuable advice, Susana Guix for providing the HAStV4 isolate, and Ian Brierley for critical reading of the manuscript. The pAVIC1 infectious clone originally developed by Suzanne Matsui was provided by Stacey Schultz-Cherry. This work was supported by Wellcome Trust grant [106207] and European Research Council grant [646891] to A.E.F.

## AUTHOR CONTRIBUTIONS

A.E.F. and V.L. conceived the project. V.L. performed the experiments. A.E.F. performed the bioinformatic analyses. V.L. and A.E.F. wrote the manuscript.

