## Supplementary for "A hidden gene in astroviruses encodes a cell-permeabilizing protein involved in virus release"

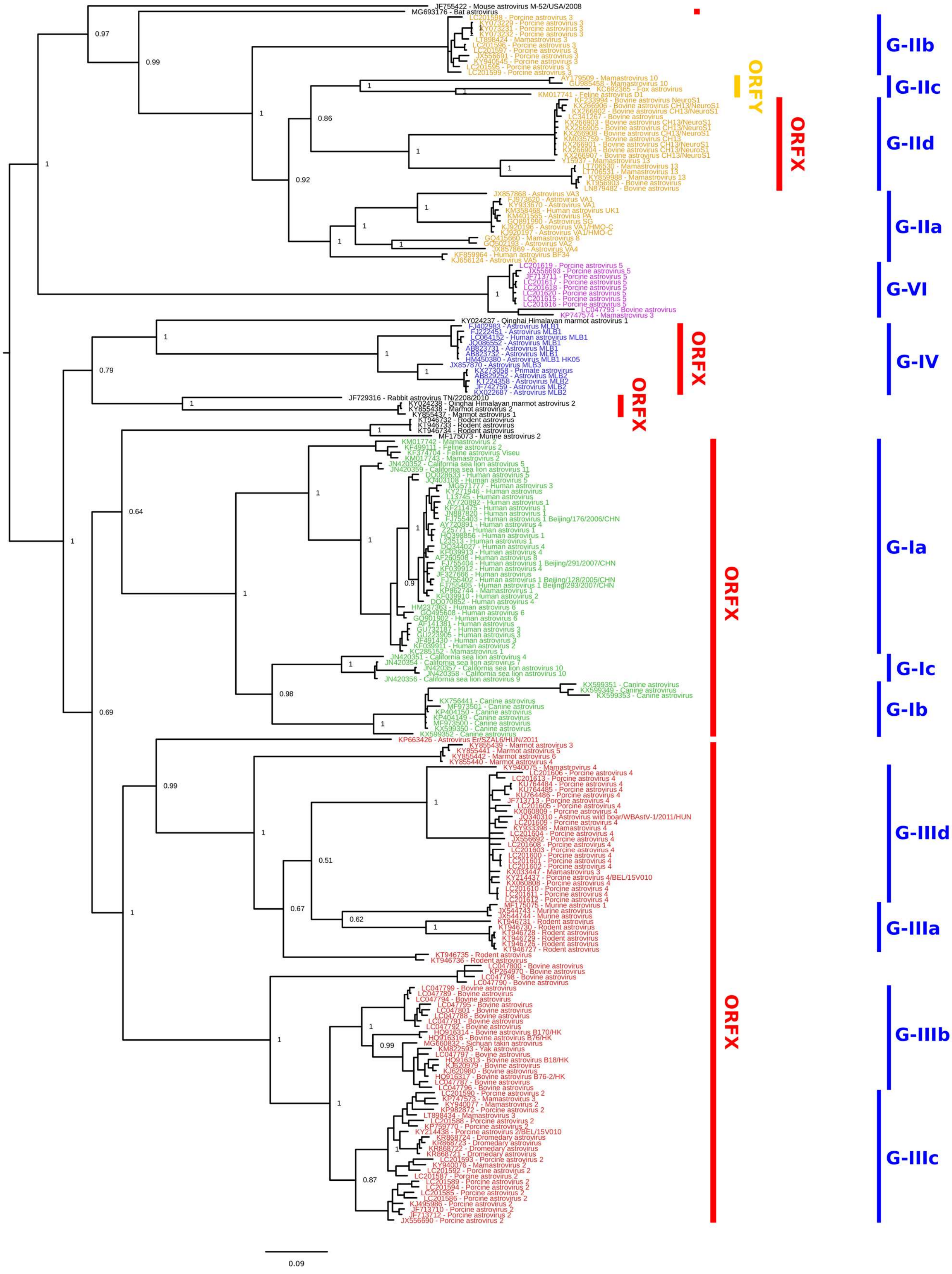


**Supplementary Figure 1 │ Phylogenetic tree of mammalian astrovirus RdRp sequences.** All full-length mammalian astrovirus genome sequences were obtained from NCBI on 26 July 2018 and the ORF1b amino acid sequences were extracted and aligned with MUSCLE [1]. A maximum likelihood phylogenetic tree was estimated using the Bayesian Markov chain Monte Carlo method implemented in MrBayes version 3.2.3 [2], sampling across the default set of fixed amino acid rate matrices, with one million generations, discarding the first 25% as burn-in. The tree was visualized with FigTree (http://tree.bio.ed.ac.uk/software/figtree/). Genogroups (based on the scheme of Yokoyama et al [3]) are indicated with coloured text: green – genogroup I, yellow – genogroup II, red – genogroup III, blue – genogroup IV, magenta – genogroup VI, unclassified sequences – black. Subgroups of sequences, defined on the basis of RdRp phylogeny for the purposes of this study only, are indicated in blue at right (G-Ia, G-Ib, etc). Taxa that have a putative ORFX (ORFY) are indicated in red (yellow) at right. The tree is midpoint rooted and deeper nodes are labelled with posterior probability values.

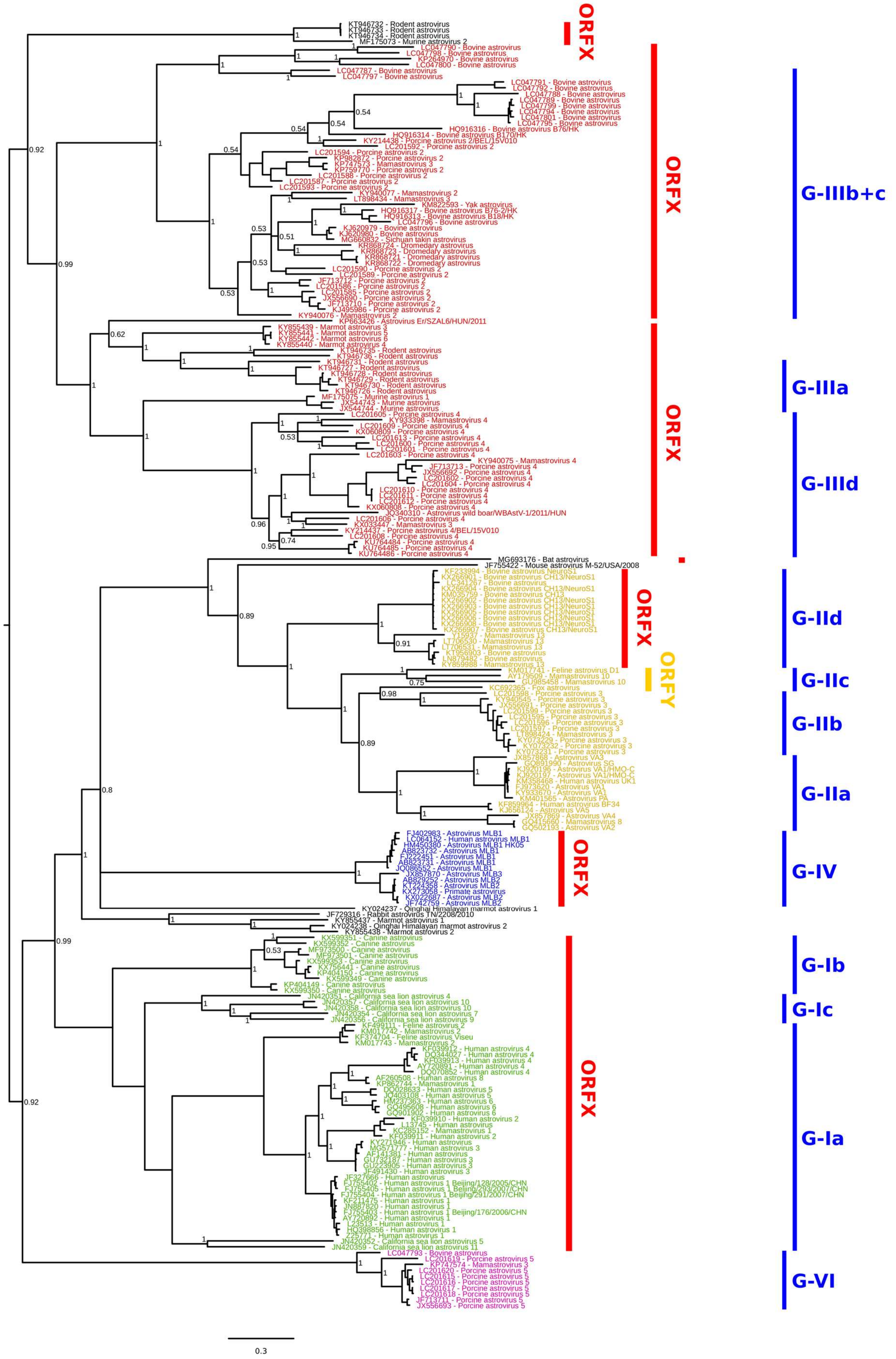


**Supplementary Figure 2 │ Phylogenetic tree of mammalian astrovirus CP sequences.** All full-length mammalian astrovirus genome sequences were obtained from NCBI on 26 July 2018 and the

ORF2 amino acid sequences were extracted and aligned with MUSCLE [1]. A maximum likelihood phylogenetic tree was estimated using the Bayesian Markov chain Monte Carlo method implemented in MrBayes version 3.2.3 [2], sampling across the default set of fixed amino acid rate matrices, with one million generations, discarding the first 25% as burn-in. The tree was visualized with FigTree (http://tree.bio.ed.ac.uk/software/figtree/). Genogroups (based on the scheme of Yokoyama et al [3]) are indicated with coloured text: green – genogroup I, yellow – genogroup II, red – genogroup III, blue – genogroup IV, magenta – genogroup VI, unclassified sequences – black. Subgroups of sequences, defined on the basis of RdRp phylogeny (Fig. S1) for the purposes of this study only, are indicated in blue at right (G-Ia, G-Ib, etc). Taxa that have a putative ORFX (ORFY) are indicated in red (yellow) at right. The tree is midpoint rooted and deeper nodes are labelled with posterior probability values.

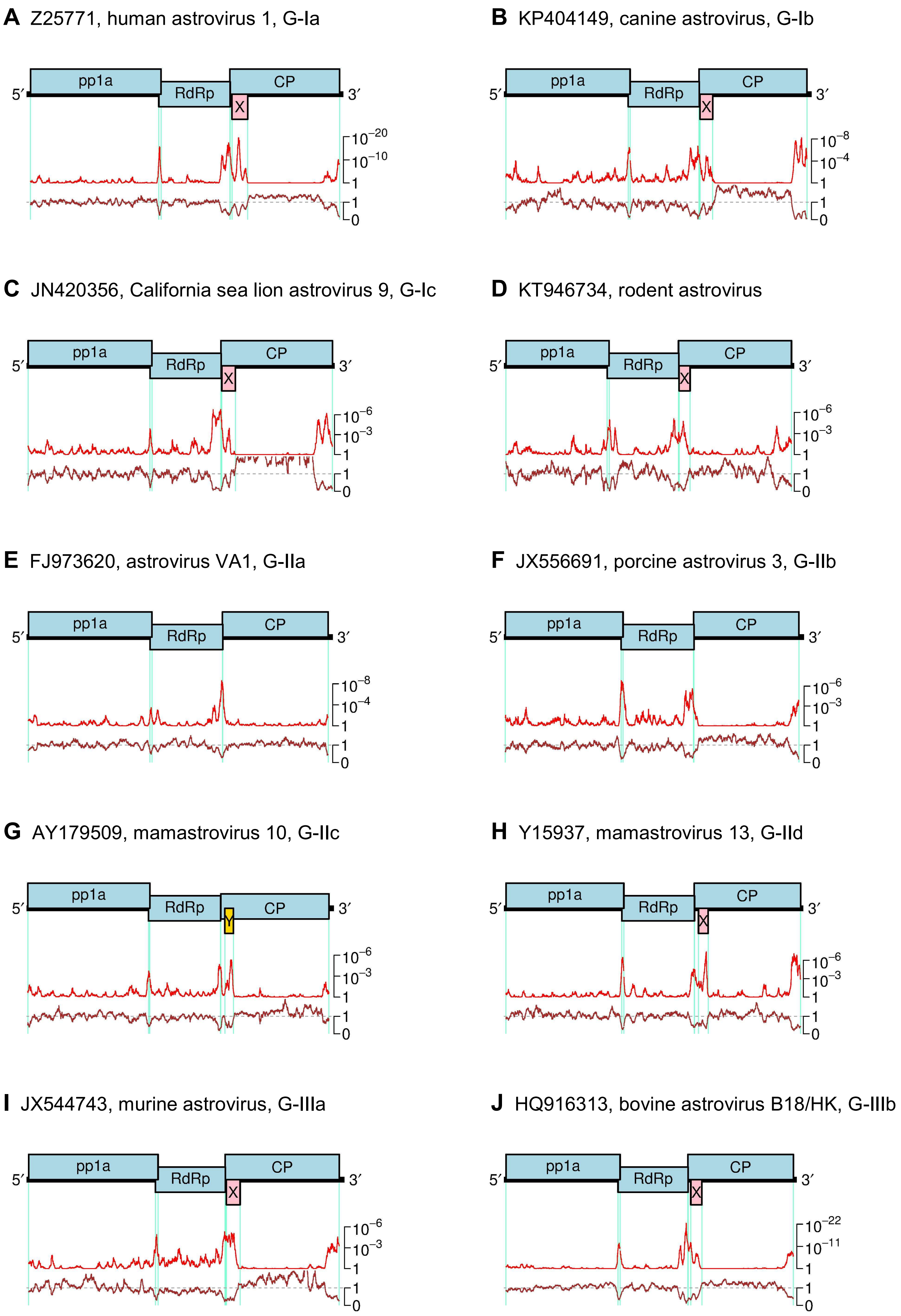
A Z25771, human astrovirus 1, G-Ia
pp1a
RdRp
CP
X
5′
3′
10−20
10−10
1
1
0
B KP404149, canine astrovirus, G-Ib
pp1a
RdRp
CP
X
5′
3′
10−8
10−4
1
1
0
C JN420356, California sea lion astrovirus 9, G-Ic
pp1a
RdRp
CP
X
5′
3′
10−6
10−3
1
1
0
D KT946734, rodent astrovirus
pp1a
RdRp
CP
X
5′
3′
10−6
10−3
1
1
0
E FJ973620, astrovirus VA1, G-IIa
pp1a
RdRp
CP
5′
3′
10−8
10−4
1
1
0
F JX556691, porcine astrovirus 3, G-IIb
pp1a
RdRp
CP
5′
3′
10−6
10−3
1
1
0
G AY179509, mamastrovirus 10, G-IIc
pp1a
RdRp
CP
Y
5′
3′
10−6
10−3
1
1
0
H Y15937, mamastrovirus 13, G-IId
pp1a
RdRp
CP
X
5′
3′
10−6
10−3
1
1
0
I JX544743, murine astrovirus, G-IIIa
pp1a
RdRp
CP
X
5′
3′
10−6
10−3
1
1
0
J HQ916313, bovine astrovirus B18/HK, G-IIIb
pp1a
RdRp
CP
X
5′
3′
10−22
10−11
1
1
0

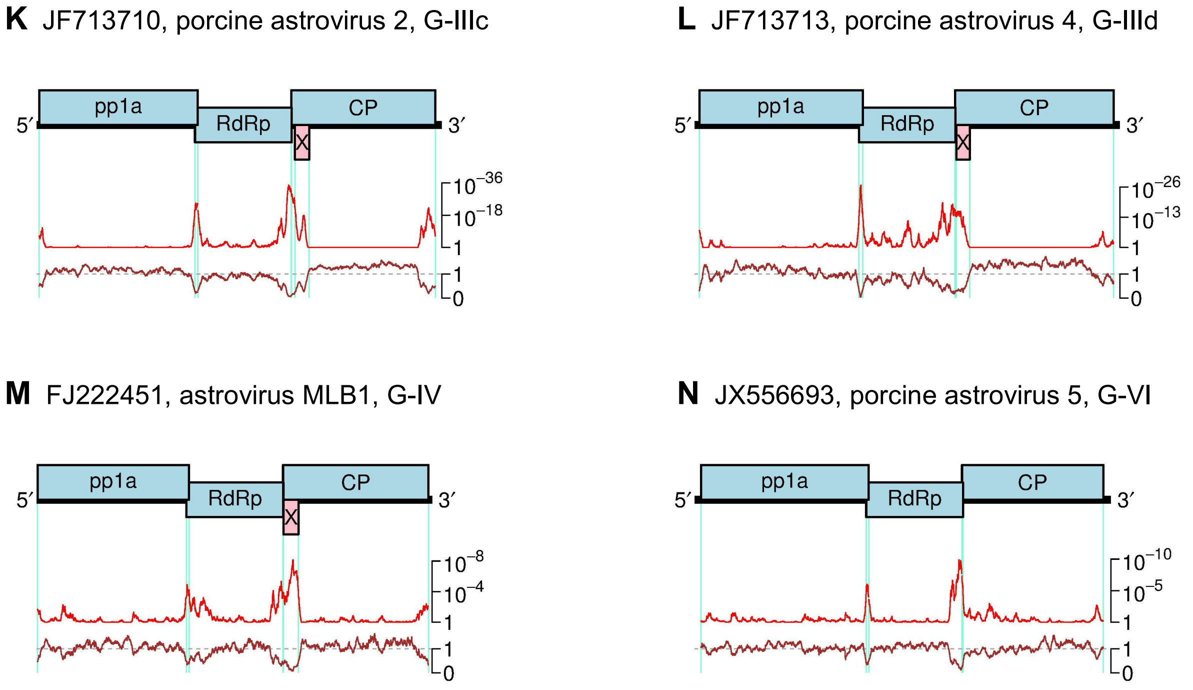


**Supplementary Figure 3 | Synonymous site conservation analysis of astroviruses.** In each subfigure, a genome map is shown at top, indicating the pp1a, RdRp and CP ORFs (blue) and the putative additional ORF where present (pink – ORFX; yellow – ORFY). Below, is shown the analysis of conservation at synonymous sites in the pp1a, RdRp and CP ORFs. The red line shows the probability that the observed conservation could occur under a null model of neutral evolution at synonymous sites, whereas the brown line depicts the ratio of the observed number of substitutions to the number expected under the null model. Peaks in synonymous site conservation may indicate functionally important overlapping elements such as the −1 PRF signal between the pp1a and RdRp ORFs, sgRNA promoter sequences, and overlapping coding sequences (i.e. the putative X and Y ORFs). Each synonymous site conservation analysis is based on an alignment of virus sequences in the indicated clade (Ia, Ib, Ic, etc) using the genome coordinate system of the indicated "reference" sequence / isolate name. The subgroup designations are defined in Fig. 1b and the sequences used in each alignment are as shown in Fig. S9.

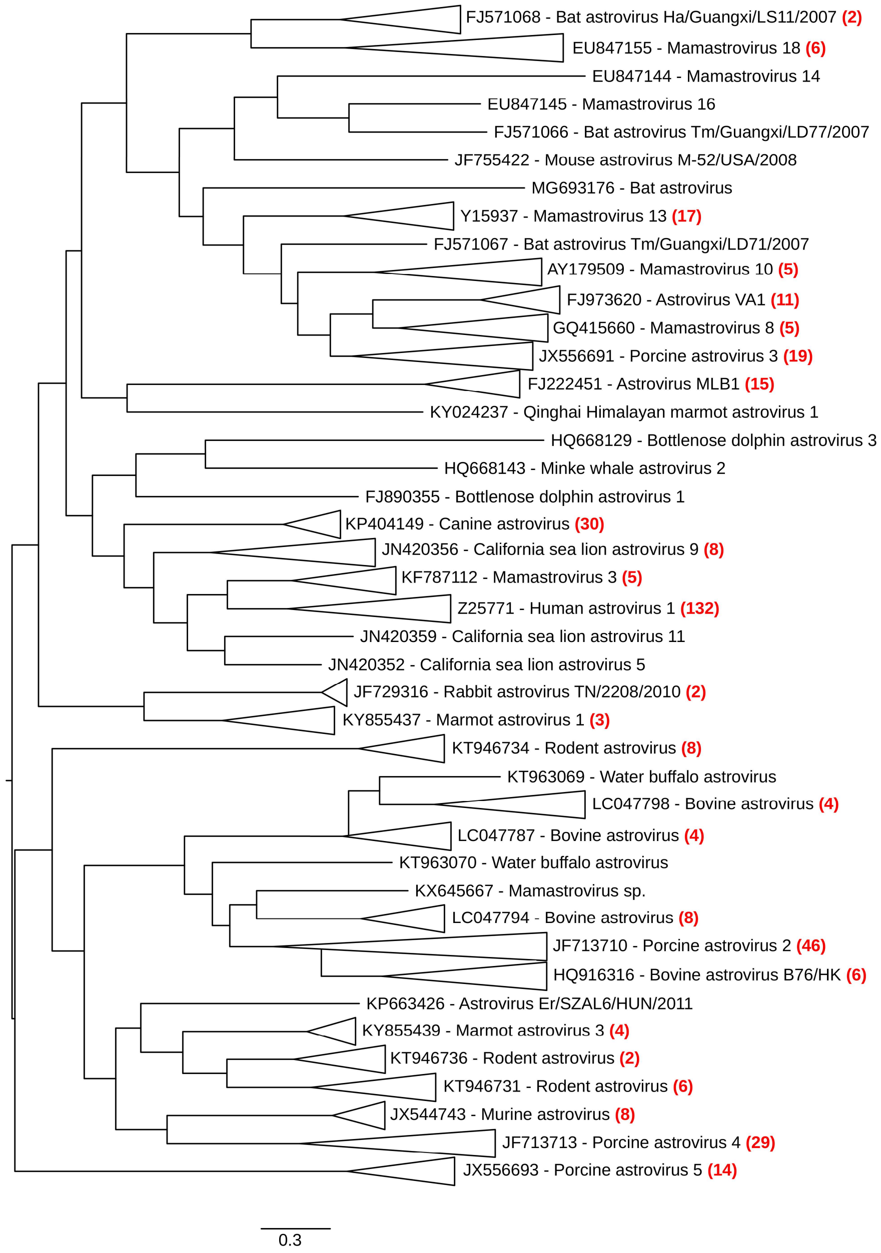


**Supplementary Figure 4 | Phylogenetic tree of mammalian astrovirus CP sequences.** Mammalian astrovirus sequences with complete or nearly complete coverage of ORF2 were obtained from NCBI on 26 July 2018, and the ORF2 amino acid sequences were extracted and aligned with MUSCLE [1]. A maximum likelihood phylogenetic tree was estimated using the Bayesian Markov chain Monte Carlo method implemented in MrBayes version 3.2.3 [2], sampling across the default set of fixed amino acid rate matrices, with five million generations, discarding the first 25% as burn-in. The tree was midpoint rooted and visualized with FigTree (http://tree.bio.ed.ac.uk/software/figtree/). Related groups of sequences (indicated by isosceles triangles) have been replaced in the figure by a single representative accession number and virus name; the total number of sequences in each group is shown in red. The 26 groups and 16 singletons correspond to those used in Fig. S5 and Fig. S7, respectively. The complete list of 415 sequences is shown in Supplementary Dataset 1.

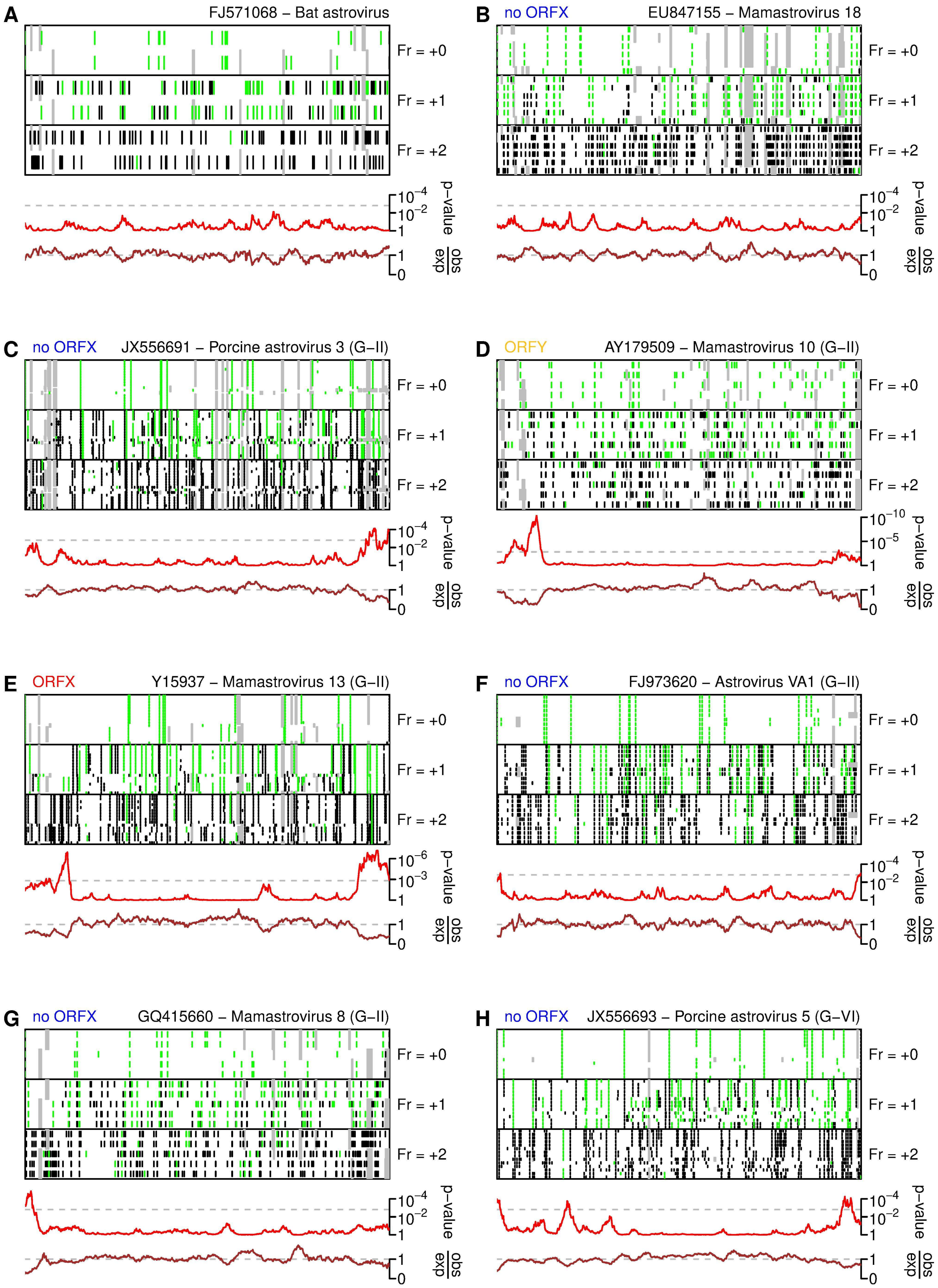

A
FJ571068 – Bat astrovirus
B
no ORFX
EU847155 – Mamastrovirus 18
C
no ORFX
JX556691 – Porcine astrovirus 3 (G–II)
D
ORFY
AY179509 – Mamastrovirus 10 (G–II)
E
ORFX
Y15937 – Mamastrovirus 13 (G–II)
F
no ORFX
FJ973620 – Astrovirus VA1 (G–II)
G
no ORFX
GQ415660 – Mamastrovirus 8 (G–II)
H
no ORFX
JX556693 – Porcine astrovirus 5 (G–VI)
Fr = +0
Fr = +1
Fr = +2
p–value
obs/exp

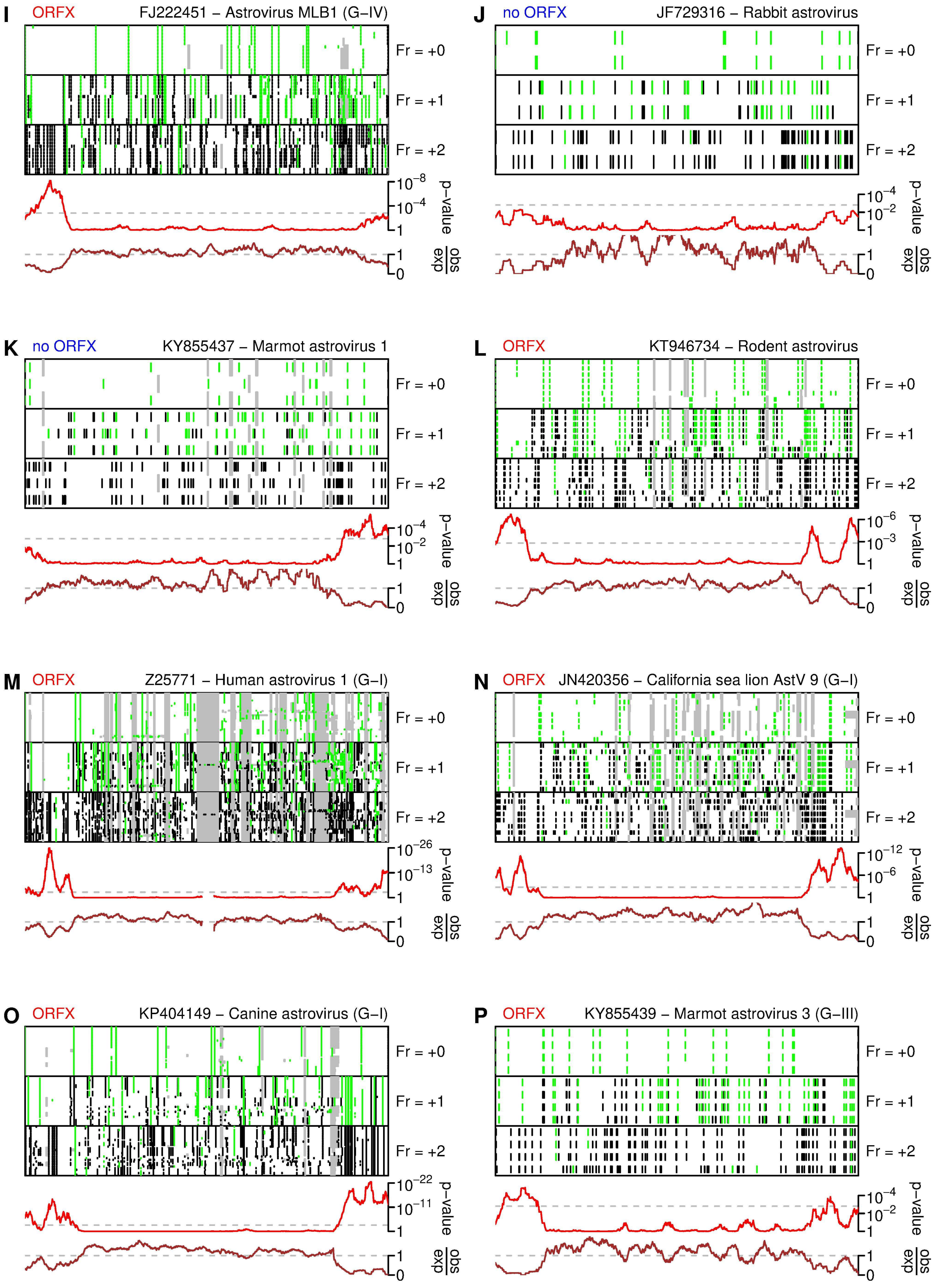
I
ORFX
FJ222451 – Astrovirus MLB1 (G–IV)
J
no ORFX
JF729316 – Rabbit astrovirus
K
no ORFX
KY855437 – Marmot astrovirus 1
L
ORFX
KT946734 – Rodent astrovirus
M
ORFX
Z25771 – Human astrovirus 1 (G–I)
N
ORFX
JN420356 – California sea lion AstV 9 (G–I)
O
ORFX
KP404149 – Canine astrovirus (G–I)
P
ORFX
KY855439 – Marmot astrovirus 3 (G–III)
Fr = +0
Fr = +1
Fr = +2
p–value
obs/exp

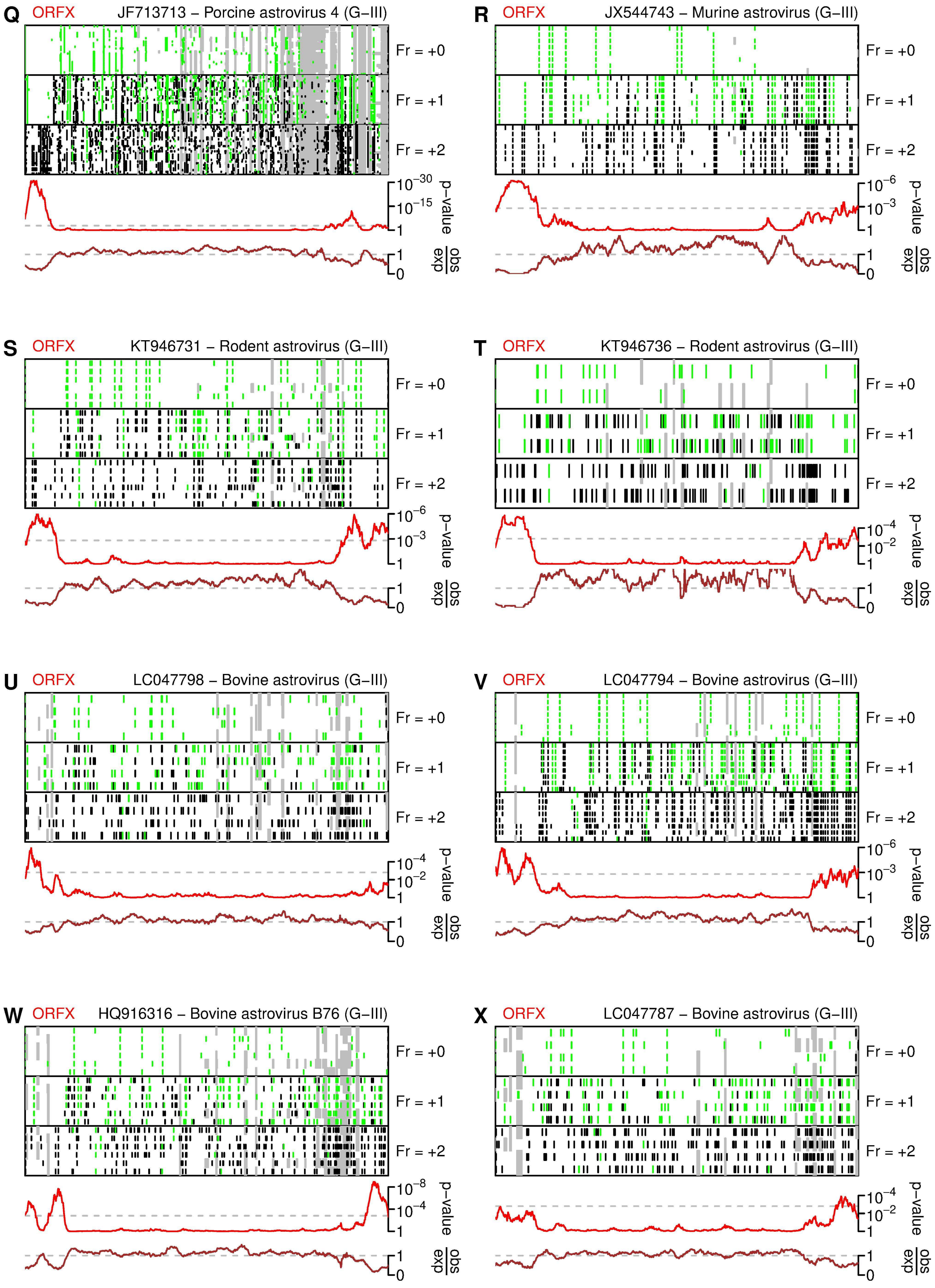
Q
ORFX
JF713713 – Porcine astrovirus 4 (G–III)
Fr = +0
Fr = +1
Fr = +2
p–value
obs/exp
R
ORFX
JX544743 – Murine astrovirus (G–III)
Fr = +0
Fr = +1
Fr = +2
p–value
obs/exp
S
ORFX
KT946731 – Rodent astrovirus (G–III)
Fr = +0
Fr = +1
Fr = +2
p–value
obs/exp
T
ORFX
KT946736 – Rodent astrovirus (G–III)
Fr = +0
Fr = +1
Fr = +2
p–value
obs/exp
U
ORFX
LC047798 – Bovine astrovirus (G–III)
Fr = +0
Fr = +1
Fr = +2
p–value
obs/exp
V
ORFX
LC047794 – Bovine astrovirus (G–III)
Fr = +0
Fr = +1
Fr = +2
p–value
obs/exp
W
ORFX
HQ916316 – Bovine astrovirus B76 (G–III)
Fr = +0
Fr = +1
Fr = +2
p–value
obs/exp
X
ORFX
LC047787 – Bovine astrovirus (G–III)
Fr = +0
Fr = +1
Fr = +2
p–value
obs/exp

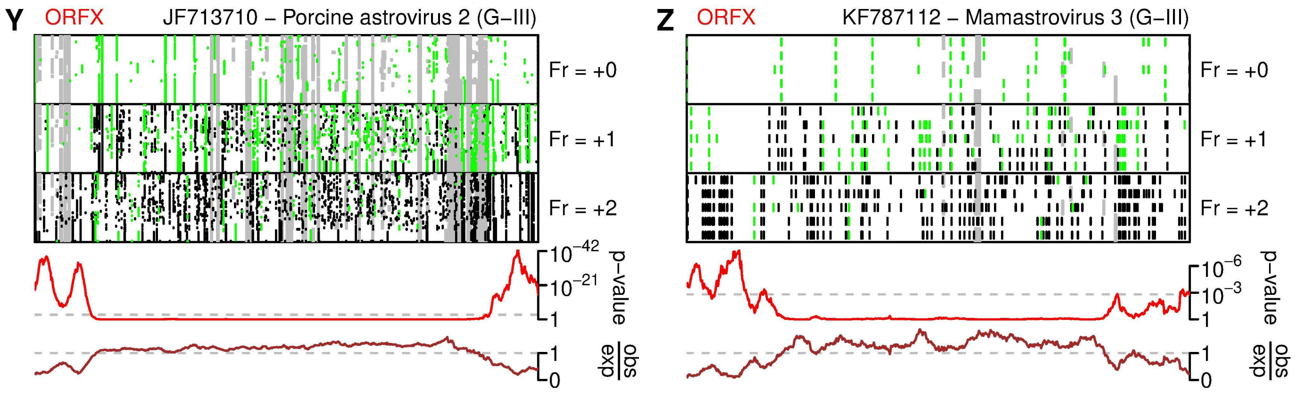


**Supplementary Figure 5 | Comparative genomic analysis of CP alignments.** Sequences with coverage of the capsid coding region of mammalian astroviruses were obtained from NCBI, CP ORF sequences were extracted, and clustered into 26 multi-sequence and 16 singleton groups based on CP amino acid identity (see Methods; Fig. S4). For each multi-sequence group, duplicate sequences were removed and remaining sequences were aligned with MUSCLE [1] and analysed with synplot2 [4]. The header of each plot indicates one sequence (accession number and virus name) from the group, genogroup (G-I, G-II etc; where defined), and whether the group is predicted to contain ORFX, ORFY or neither. The upper three panels show the positions of alignment gaps (grey), stop codons (black) and AUG codons (green) in each of the three reading frames in each sequence in the alignment. Below, is shown the analysis of conservation at synonymous sites. The red line shows the probability that the observed conservation could occur under a null model of neutral evolution at synonymous sites, whereas the brown line depicts the ratio of the observed number of substitutions to the number expected under the null model. Peaks in synonymous site conservation may indicate functionally important overlapping elements such as overlapping coding sequences (i.e. the putative X and Y ORFs) besides regulatory elements (e.g. functional RNA structures). See Fig. S7 for the singleton groups.

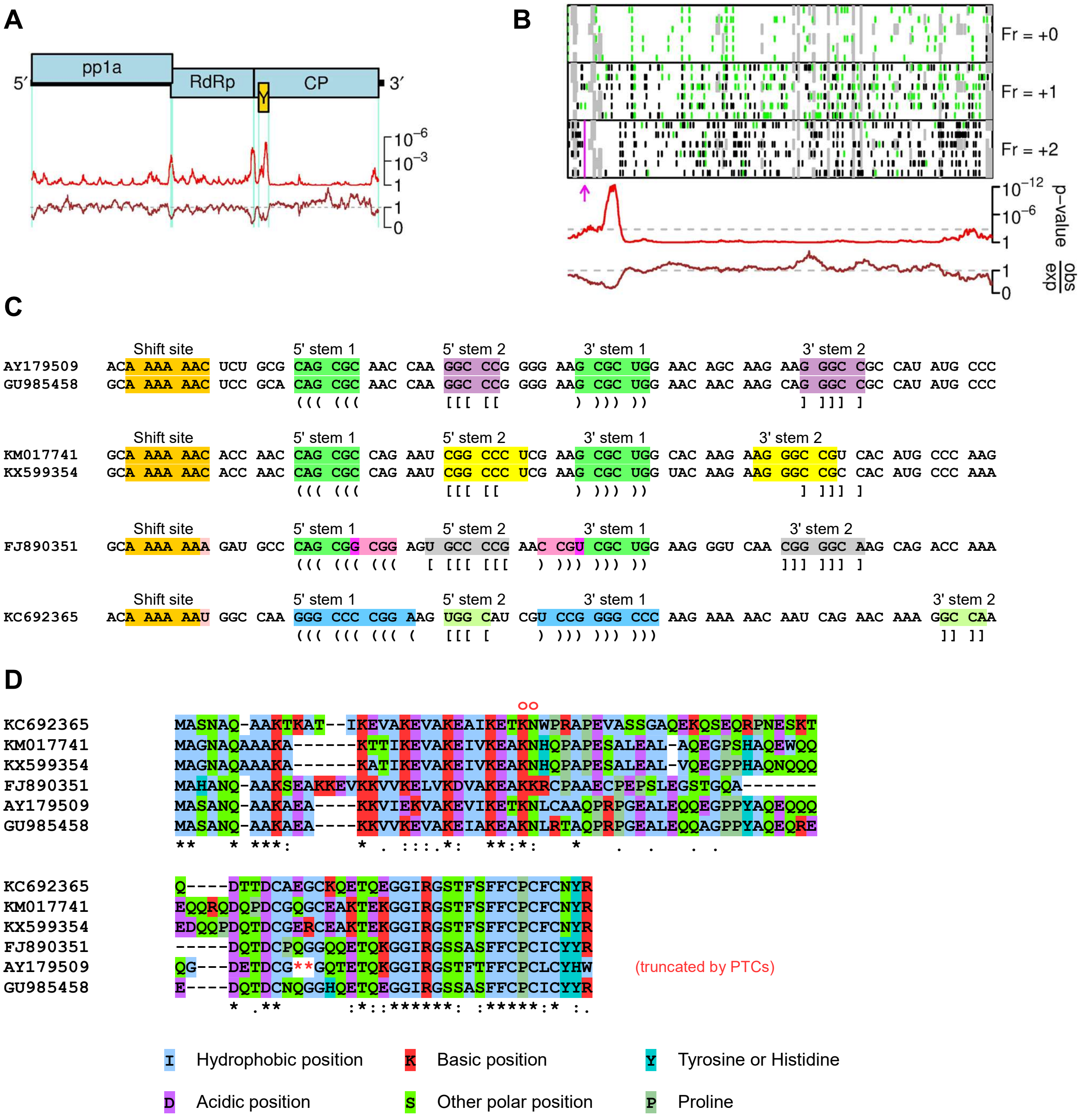


**Supplementary Figure 6 | A putative −1 PRF overlapping ORF in subgroup IIc astroviruses. (A)** Comparative genomic analysis of subgroup IIb full-length sequences AY179509, GU985458, KC692365 and KM017741. Top – genome map indicating the pp1a, RdRp and CP ORFs (blue) and the putative additional Y ORF (yellow). Bottom – analysis of conservation at synonymous sites in the pp1a, RdRp and CP ORFs. The red line shows the probability that the observed conservation could occur under a null model of neutral evolution at synonymous sites, whereas the brown line depicts the ratio of the observed number of substitutions to the number expected under the null model. Peaks in synonymous site conservation may indicate functionally important overlapping elements such as the −1 ribosomal frameshifting signal between the pp1a and RdRp ORFs, sgRNA promoter sequences, and overlapping coding sequences (i.e. the putative Y ORF). **(B)** Synonymous site conservation plot for the CP ORF for AY179509, GU985458, KC692365, KM017741 and two additional partial-genome sequences with CP ORF coverage, FJ890351 and KX599354 (lower two panels). In the upper three panels, alignment gaps (grey), stop codons (black) and AUG codons (green) are indicated in each of the three reading frames. The location of the putative A_AAA_AAZ −1 ribosomal frameshifting site is indicated in pink. **(C)** Predicted −1 PRF shift sites and 3′-adjacent

RNA secondary structures in each of the six sequences. Predicted base-pairings are indicated with matching highlights and "()"s (stem 1) or "[]"s (stem 2). **(D)** Alignment of the predicted −1 PRF product. The location of the A_AAA_AAN shift site is indicated with two red circles. The upstream sequence is encoded by the 5′ end of the CP ORF whereas the downstream sequence is encoded by the overlapping −1 frame Y ORF. Amino acids are colour-coded according to their physicochemical properties. In AY179509, the Y ORF is truncated by two tandem premature termination codons (red asterisks).

**EU847145 - Mamastrovirus 16**

MRLEINLTKRISTRVSKSMLGRRLRRKRGRMVMVDLNLNLLLMLLLLLGLLRAVMSLL
????????????????????????

**FJ890355 - Bottlenose dolphin astrovirus 1**

MLALRLKPPAHREAKVVPVQGLEEEHPLSKSQLIPKQKDLPEDQVDALLELKIIVSNNKLEINSRNKVSQGPPQRLSRPRRLLLALLDQILAMMQRGRFPSI
??????????????????

**HQ668129 - Bottlenose dolphin astrovirus 3**

MAAKSQLKSSPEPELRLVKAITRELVEEEKEVIRTIKRMLANNQEIIITRLGVSEQRYHVQLTAWAYADPNQKIKSSSLQCLVPSVETHQEGSKWRRWFSLTRCSLRKSQGITT

LDHCK

**HQ668143 - Minke whale astrovirus 2**

MPHNLKRNASVSGVTQEELELLEILTAILESHDEVIVDVINRMEGRLELRLNAWDSEGLSQHSHRLSRPRWAPLVPMRRLFLRLKGFFTSTPHSLRRSPGVWRLDRFRP

**JN420352 - California sea lion astrovirus 5**

MAATGANLVPAPRVGAGNQMSRSQSIQNPEEQEVELDAVNVNLITVSVNLSNNSLTSLVLQDQSQPSDKKPLLRLALSTLIRLIRRSLRR
?????????????????

**JN420359 - California sea lion astrovirus 11**

MKNPTGAILAAAHNPEAATRQSRSPSIQRSQEVETDAVNVNLVSVSVQLSINNSGNRVSQDQKCQLNSGQQQHSEPLEPTPLGSQSLRRASFVTRFLSRTVQVLRLLAQFRC

**KT963069 - Water buffalo astrovirus**

MAMGGMPRAWWFRPLRVTNRHRTKVDGVDGARLRSMSTSQTRQDHQEDQQDNNNNLRDNVNFGCGATAALRAVKLRFSIRYARR
??????????????????

**KT963070 - Water buffalo astrovirus**

MDQHPQEQVDDVDGVEPNPENPKLLCCLPLRKFRFRVSGPAVRASITMWFGRESPPPLEQLEPTPTGK

**KX645667 - Mamastrovirus sp.**

MEDQHNPPPNPSRDETDGRDIMRRLRRLYALYKELKNVADQILQEDVVEGGLYCRKLQQRSEQLDQMLQKQSKTSSQSSSIPPP

**MG693176 - Bat astrovirus**

MPSLRTRRLRLRSRLNLISNILRNTNRNRPRARVRLEQSLRRLRRTLRRSVLRGRNRGRQSVSRPLSASLTGPPRRARCWLLVQIFILLLPRNLKVALLSALWLRRRPSLANGA

FLGLLSVLPPWLALLLLLALSLAFPSIQQAGPFQ

**Supplementary Figure 7 | Potential XP sequences in unclustered astroviruses.** Sequences with coverage of the capsid coding region of mammalian astroviruses were obtained from NCBI, CP ORF sequences were extracted, and clustered into 26 multi-sequence and 16 singleton groups based on CP amino acid identity (see Methods). Potential XP peptides encoded within 10 of the 16 unclustered sequences are shown below. Amino acids are colour-coded according to their physicochemical properties. Transmembrane regions predicted by Phobius [5] are indicated with pink bars. Potential transmembrane regions (hydrophobic regions scored below threshold by Phobius) are indicated with pink question marks. See Fig. S5 for the multi-sequence groups.

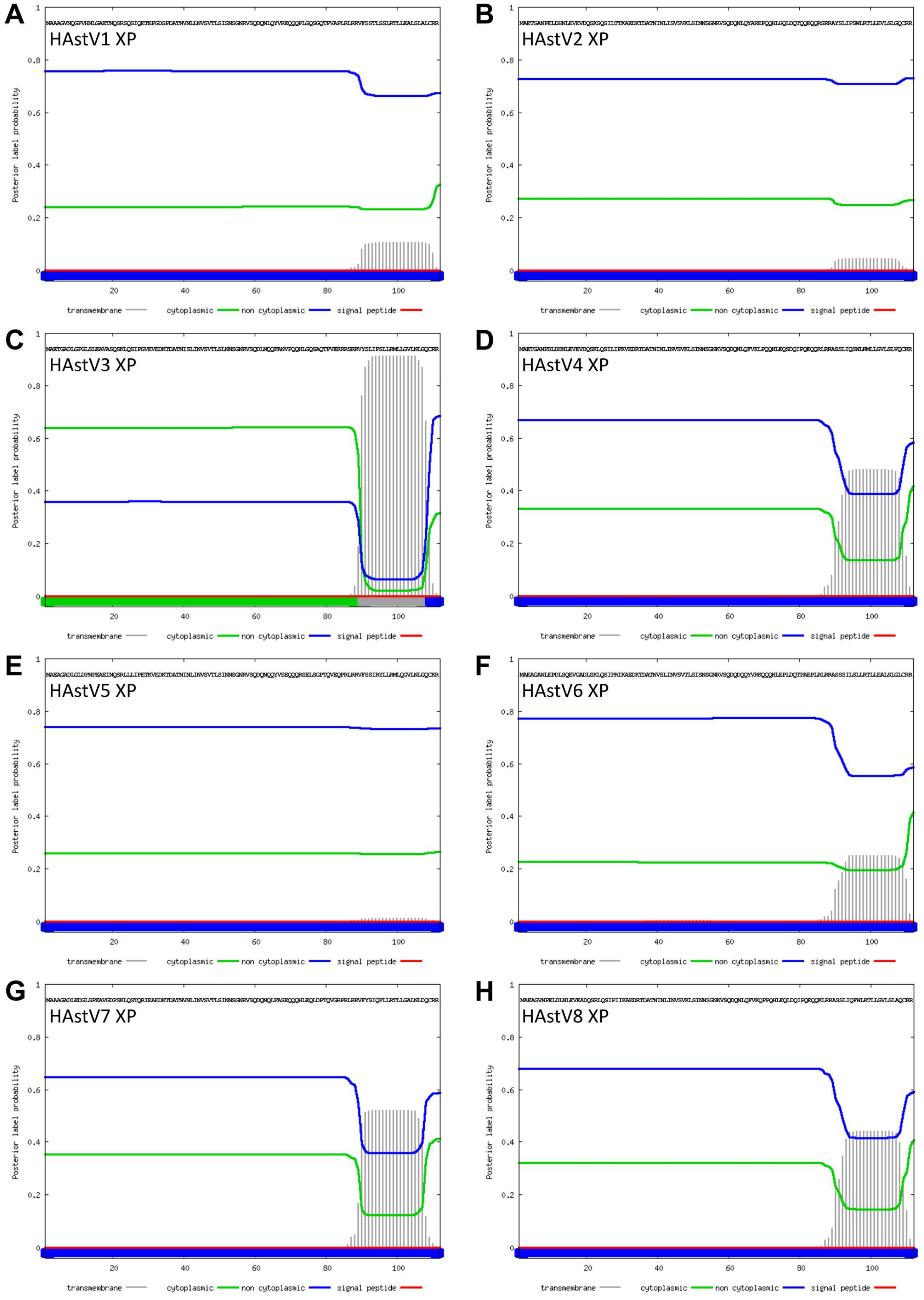


**Supplementary Figure 8 | Transmembrane domain predictions for HAstV XPs.** Transmembrane domains for representative sequences (see Fig. 1F) were predicted with Phobius [5]. An above-threshold TM was predicted only for HAstV3.

I Hydrophobic position

K Basic position

Y Tyrosine or Histidine

D Acidic position

S Other polar position

P Proline

**Bat astrovirus**

```
MG693176 MPSLRTRRLRLRSRLNLISNILRNTNRNRPRARVRLEQSLRRLRRTLRRSVLRGRNRGRQSVSRPLSASLTGPPRRARCWLLVQIFILLLPRNLKVALLSALWLRRRPSLANGAFLGLLSVLPP
         WLALLLLLALSLAFPSIQQAGPFQ
```

**Genogroup IId - bovine astrovirus etc**

```
KF233994 MLHLPSRLPRRLLNLRRRQKINNNRPSARPSARTGFRVEVETRQVKREFQIELKQSSKSRVWKVLTLGLGSLCLLLLVRLVLTSPRALSYR
KX266906 MLHLPSRLPRRLLNLRRKQRTNNNRPSARPSARTGFRVEVETRQVKREFQIELKQSSRSKVWKALTLGLGLLFLLLLVRLVLISPRALSYR
KX266902 MLHLPSRLPRRLLNLRRKQRTNNNRPSARPSARTGFRVEVETRQVKREFQIELKQSSRSKVWKALTLGLGLLFLLLLVRLVLISPRALSYR
LC341267 MLHLPSRLPRRLLNLRRRQKISNNRPSARPSARTGFRVEVETKQVKREFQIELKQSSKSRVWKVLTLGLGLLCLLLLVRLVLTSPRALSYR
KX266903 MLHLPSRLPRRLLNLRRKQRTNNNRPSARPSARTGFRVEVETRQVKRGFQIELKQSSRSKVWKVLTLGLGLLFLLLLVRLVLTSPRALSYR
KX266905 MLHLPSRLPRRLLNLRRKQRTNNNRPSARPSARTGFRVEVETRQVKREFQIELKQSSRSKVWKVLTLGLGLLFLLLLVRLVLTSPRALSYR
KX266908 MLHLPSRLPRRLLNLRRKQRTNNNRPSARPSARTGFRVEVETRQVKREFQIELKQSSRSKVWKVLTLGLGLLFLLLLVRLVLTSPRALSYR
KM035759 MLHLPSRLPRRLLNLRRKQRTNNNRPSARPSARTGFRVEVETRQVKREFQIELKQSSRSKAWKVLTLGLGLLFLLLLVRLVLTSPRALSYR
KX266901 MLHLPSRLPRRLLNLRRKQRTNNNRPSARPSARTGFRVEVETRQVKREFQIELKQSSRSKVWKVLTLGLGLLFLLLLVRLVLTSPRALSYR
KX266904 MLHLPSRLPRRLLNLRRKQRTNNNRPSARPSARTGFRVEVETKQVKREFQIELKQSSRSKVWKVLTLGLGLLFLLLLVRLVLTSPRALSYR
KX266907 MLHLPSRLPRRLLNLRRKQRTNNNRPSARPSARTGFRVEVETRQVKREFQIELKQSSRNKVWKVLTLGLGLLFLLLLVRLVLTSPRALSYR
Y15937   ---------MGSSILRRM-----SPRASGGLARLSLTIGVSLTRSRG--------NYTNRGLKVLLLGLGLLSLLLLARLALIRNRDLSYR
LT706530 -------------MLRRIRSNINSSLAVIGRPSLDLRLDLLTLRLRR--------SSTSRGLRALQLGLGLMFLLLLARLALMPNKVQSCR
LT706531 -------------MLRRIRSNINSSLAVIGRPSLDLRLDLLTLRLRR--------SSTSRGLRALQLGLGLMFLLLLARLALMPNKVQSCR
KY859988 -------------MLRRIRTNINSKLAVIGRPNLDLRLDLLTLRLRR--------SSTSRGLRALQLGLGLMFLLLLARLALMPSKAQSCR
KT956903 -------------MLRRIRTNINSKLAVIGRPNLDLRLDLLTLRLRR--------SSTSRGLRALQLGLGLMFLLLLARLALMPSKAQSCR
LN879482 -------------MLRRIRTNISSKLAVIGRPNLDLRLDSLTLRLRR--------NSTSRGLRALQLGLGLMFLLLLARLALMPSKAQSCR
```

**Genogroup IV - MLB astroviruses**

```
FJ402983 MPVKVLQSTSTMQNGSLALPITSELELLGQILHQHPNLGNVGIFLIGIAVVGKILVQLGLNLRCRRQSQQHSAPLDQI
FJ222451 MPVKVLQSTSTMQNGSLASPITSELDLLGQILHQHPNLGNVGLFLIGIAVVGKILVQLGLNLRCRRQLQQHSAPLDQI
LC064152 MPVKVLQLTSTMQNGSLASPITSELDLLGQILHQHPNLGNVGIFLIGIAVVGKILVQLGLNLRCRRQLQQHSAPLDQI
JQ086552 MPVKVLQLTSTMQNGSLASPITSELDLLGQILHQHPNLGNVGIFLIGIAVVGKILVQLGLNLRCRRQLQQHSAPLDQI
AB823731 MPVKVLQLTSTMQNGSLASPITSELDLLGQILHQHPNLGNVGLFLIGIAVVGKILVQLGLNLRCRRQLQQHSAPLDQI
AB823732 MPVKVLQLTSTMQNGSLASPITSELDLLGQILHQHPNLGNVGLFLIGIAVVGKILVQLGLNLRCRRQLQQHSAPLDQI
HM450380 MPVKVLQLTSTMQNGCLASPITSELDLLGQILHQHPNLGNVGLFLIGIAVVGKILVQLGLNLRCRRQLQQHSAPLDQI
JX857870 -----------MSNASNVLQVTSELNLLDLVLSEHSNLGNVGIFLIGIAVVGRILIQLGLNLRCRKQSLQHSAPLDQI
KX273058 -----------MSNVSSVLPVTSELDLLGQVLSEHPSLGNVGIFLIGIAVVGRILIQLGLSLRCHRRSQQHLAPLDQI
AB829252 -----------MSNVSSVLPVTSELDLLGQVLSEHPSLGNVGIFLIGIAVVGRILIQLGLSLRCHRRSQQHLAPLDQI
KT224358 -----------MSNVSSVLPVTSELDLLGQVLSEHPSLGNVGIFLIGIAVVGRILIQLGLSLRCHRRSQQHLAPLDQI
JF742759 -----------MSNVSSVLPVTSELDLLGQVLSEHPSLGNVGIFLIGIAVVGRILIQLGLSLRCHRRSQQHLAPLDQI
KX022687 -----------MSNVSSVLPVTSELDLLGQVLSEHPSLGNVGIFLIGIAVVGRILIQLGLSLRCHRQSQQHLAPLDQI
```

**Rodent astrovirus**

```
         sssssssssssssssssssssss
KT946732 MLRDPVALALVLLWVVTLGAVVAALVNVGLKDPTASALCAASLLQWAVVGHALHARLEQLQDKCSGERDQEPLLANGQP
KT946733 MLRDPVALALVLLWVVTLGAVVAALVNVGLKDPTASALCAASLLQWAVVGHALHARLEQLQDKCSGERDQEPLLANGQP
KT946734 MLRDPVALALVLLWVVTLGAVVAALVNVGLKDPTASALCAASLLQWAVVGHALHARLEQLQDKCSGERDQEPLLANGQP
MF175073 MLRDLVALALAGLWVLTLGAVVAALVNVGLKDPVAASLCASSLIMWAMVGHALHARLERLPDGCSEMKDHERLLGKDLP
```

**Genogroup Ia - human astrovirus etc**

```
                                                                                                            ?????????????????????
KM017742 -MAEAGVDLDPDPDLGVEAKQSRLQLIPRQVVKGDKTDGANVSLLNVSVVLSINNSGNRASQDLNQQYVREQQLHSGPSDQTPVEQLRSRRVFSSTLSLLRMLLEALSLGLCRR
KF499111 -MAEAGVSPDPVPNLGVEAKQLRLQLTPEAVAKEDRTDGANVNLLNVSVTLSINNSGNRVSQDLNQQYVRRQQLPSGQLDQTPVEQPRLKRAYSSTLSLLRMLLEALSLGLCKR
KF374704 -MAEAGASPGPALDLGDEAKQLRLRSIPEVAAKEDKTDGANVNLLNVSVTLSINNSGNRVSQDLNQQYVREQQLHSGQLDQTPAEQLRSRRAYSSTLSLLRMLLEVLSLGLCRR
KM017743 -MAEAGVSPDPAPNLGAEAKQLKLRLTPEAVAREDKADGANVSLLNVSVTLSINNSGNRVSQDQNQQYVRRQQLHSGPSDQTPVEQLRSRRVFSSTLSLLRMLLEALSLGLCRR
JN420352 -MAATGANLVPAPRVGAGNQMSRSQSIQNPEEQEVELDAVNVNLITVSVNLSNNSLTSLVLQDQSQ-----------PSDKKPL----LRLALSTLIRLIRRSL--------RR
JN420359 MKNPTGAILAAAHNPEAATRQSRSPSIQRSQ--EVETDAVNVNLVSVSVQLSINNSGNRVSQDQKCQLNSGQQQHSEPLEPTPLGSQSLRRASFVTRFLSRTVQVLRLLAQFRC
DQ028633 -MAEAGADLGLDPNPEAEINQSRLLLIPETKV-EDKTDATNINLINVSVTLSINNSGNRVSQDQNQQYVSEQQQRSELSGPTQVEQPRLRRVFSSIRYLLRMLQGVLNLGQCKR
JQ403108 -MAGAGADLGLDLNPEAETNQSKLQLIPETKV-EDKTDATNINLINVSVTLSINNSGNRVSQDQNQQYVNEQQQRSELSDPIQVEQPRLRRVFSSIRYLLRMLQGVLNLDQCRR
MG571777 -MAETGADLGPGLSLEAVANQSKLQLTPGVEA-EDKTDATNISLINVSVTLSLNNSGNRVSRDLNQQFVNVPQQHSGQSARTPVERRRSRRVYSLIPSLLRMLLGVLNLGQCRR
KY271946 -MAETGADLGPGLSLEAVANQSKLQLTPGVEA-EDKTDATNISLINVSVTLSLNNSGNRVSRDLNQQFVNVPQQHSGQSAQTPVERRRSRRVYSLIPSLLRMLLGVLNLGQCRR
L13745   -MAETGANPELDHNLEVEVDQSKSQSILTTKA-EDKTDATNINLISVSVKLSINNSGNRVSQDQNLQYAREPQQHLGQLDQTQQEQQRSRRAYSLIPSWLRTLLEVLSLGQCRR
AY720892 -MAATGVNQGPDHNLGAETNQSRLQSIQEAEL-GDSPDATSVNLLNVSVTLSINNSGNKVSQDQNLRYVREQQQPLGQSGQTPVALPRLRRVFSSTLSLLRTLLGALSLALCRR
KF211475 -MVATGVNQEPVHNLGAETNQSRLQSIQEAEL-GDSPDATNVNLLNVSVTLSINNSGNRVSQDQNLRYVREQQQPLGQSGQTPVALPRLRRVFSSTLSLLRTLLGALSLALCRR
JN887820 -MAATGVNQGPVHNLGAETNQSRLQSIQEAEL-GDSPDATNVNLLNVSVTLSINNSGNKVSQDQNLRYVREQQQPLERSGQTPVALPRLRRVFSSTLSLLRTLLGALSLALCRR
FJ755403 -MAATGVNQGPVHNLGAETNQSRLQSIQEAEL-GDSPDATNVNLLNVSVTLSINNSGNKVSQDQNLRYVREQQQPLERSGQTPVALPRLRRVFSSTLSLLRTLLGALSLALCRR
AY720891 -MAEAGANPELDHNLEVEVDQSKLQSIPIIKA-EDRTDATNINLISVSVKLSINNSGNKVSQDQNLQFVKQPPQHLEQLDQTPQEQQKLRRASSLIQSWLRTLLGVLNLAQCRR
Z25771   -MAAAGVNQGPVHNLGAETNQSRSQSIQETEP-GDSPDATNVNLLNVSVTLSISNSGNRVSQDQNLQYVREQQQPLGQSGQTPVAPLRLRRVFSSTLSSLRTLLEALSLALCRR
HQ398856 -MAAAGVNQGPAHNLGAETNQSRSQSIQETEP-GDSPDATNVNLLNVSVTLSISNSGNRVSQDQNLQFVREQQQPLGQSGQTPVAPLRLRRVFSSTLSSLRTLLEALSLALCRR
L23513   -MAATGVNQGPVHNLGAEINQSRLQSIQETEP-GDSPDATNVNLLNVSVTLSISNSGNRVSQDQNLQYVREQQQPLGRSGQTPVAPLRLRRVFSSTLSSLRTLLEALSLALCRR
DQ344027 -MAEAGANPELDLNLEVEADQSRLQSIPIIKA-EDKTDATNINLINVSVKLSINNSGNKVSQDQNLQFAKQPPQHLEQLDQTPQEQQKLRRASSLIQFWLRTLLGVLSLAQCRR
KF039913 -MAEAGANPELDLNLEVEADQSRLQSIPIIKA-EDKTDATNINLINVSVKLSINNSGNKVSQDQNLQFVKQPPQHLEQLDQIPQEQQKLRRASSLIQFWLRTLLGVLSLAQCRR
AF260508 -MAEAGVNPELDLNLEVEADQSRLQSIPIIKA-EDRTDATNINLINVSVKLSINNSGNKVSQDQNLQFVKQPPQHLEQLDQIPQEQQKLRRASSLIQFWLRTLLGVLSLAQCRR
FJ755404 -MAATGANQEPARNLEGGTDQLKSQLIQKTGP-EDSPDAANINLLNVSVTLSISNSGNKVSQDQNLQYVRGQQQPLEQSGQTPVAPLRLRRVFSSTLSLLRTLLEVLSLAPCRR
KF039912 -MAEAGANPDLDLNLEVEADQSRLQSIPIIKA-EDRTDATNINLINVSVKLSINNSGNRVSQDQNLQFVKQPPQHLEQLDQIPQEQQKLRRASSLIQFWLRTLLGVLSLAQCRR
JF327666 -MAATGVNQGPAHNLEGGTEQLKSQLIQKTGL-EDSPDAANINLLNVSVTLSISNSGNKVSQDQNLQYVRGQQQPLGRSGQTPAAPLRLRRVFSSTLSLLRTLLGALSLAPCRR
FJ755402 -MAVTGVNQEPAHNLEGGTDQLKSQLIQKTGP-EDSPDAANINLLNVSVTLSISNSGNKVSQDQNLQYVRGQQQPLGRSGQTPVAPLRLRRVFSSTLSLLRTLLGALSLAPCRR
FJ755405 -MAVTGVNQEPAHNLEGGTDQLKSQLIQKTGS-EDSPDAANINLLNVSVTLSISNSGNKVSQDQNLQYVRGQQQPLGRSGQTPVAPLRLRRVFSSTLSLLRTLLGALSLAPCRR
KP862744 -MAEAGANPELDLNLEVEADQSRLQLIPITKA-EDRTDATNINLINVSVKLSINNSGNKVSQDQNLQFVKQPPQHSEQLDQIPQEQQKLRRASSLIQFWLRTLLGVLSLAQCRR
KF039910 -MAETGANLDLGHNQEVEVEQSKLQSTPRIRA-EDKMDATNISLINVSVKLSISNSGNRVSQDQNLQYASEPPQHSAQLAQTPQEQQRSRRAYFSIPSWLRMLLGVLSLDQCRH
DQ070852 -MAETGANPDLDHNLEVEVDQSKLQSILIPKV-EDRTDATNINLINVSVKLSINNSGNKVSQDQNLQFVKLPQQHLEQSDQIPQEQQRLRRASSLIQSWLRMLLGVLSLVQCRR
HM237363 -MAEAGANLEPDPSQEVGVDQSKLQSIPRIKA-EDKTDATNVSLINVSVTLSISNSGNRVSQDQNQQYVRKQQQHLGPLDQTPVEPLRLRRVSSSILSLLRTLLEALSLGLCRR
GQ495608 -MAEAGANLEPDLSQEVGADLSKLQSIPRIKA-EDKTDATNVSLINVSVTLSISNSGNRVSQDQDQQYVRKQQQHLEPLDQTPAEPLRLRRASSSILSLLRTLLEALSLGLCRR
GQ901902 -MAEAGANLEPDLSQEVGADLSKLQSTPRIKA-EDNTDATNVSLINVSVTLSISNSGNRVSQDQNQQYVRKQQQHLGPLDQTPVEPLRLRRVSSSILSLLRTLLEALSLGLCRR
AF141381 -MAETGADLGPGLSLEAVASQSKLQSIPGVEV-EDKTDATNISLINVSVTLSLNNSGNRVSQDLNQQFANVPQQHLGQSAQTPVERRRSRRVYSLIPSLLRMLLGVLNLGQCRR
GU732187 -MAETGADLGPGRSLEAVANQSKLQSIPGVEV-EDRTDATNISLINVSVTLSLNNSGNRVSQDLNQQFANVPQQHLGQSAQTPVERRRSRRVYSLIPSLLRMLLGALNLGQCRR
GU223905 -MAETGADLGPGLSLEAVASQSKLQSIPGVEV-EDKTDATNISLINVSVTLSLNNSGNRVSQDLNQQFANVPQQHLGQSAQTPVERRRSRRVYSLIPSLLRMLLGALNLGQCRR
JF491430 -MAETGADLGPGLSLEAVASQSKLQSIPGVEV-EDKTDATNISLINVSVTLSLNNSGNRVSQDLNQQFANVPQQHLGQSAQTPVERRRSRRVYSLIPSLLRMLLGALNLGQCRR
KF039911 -MAGTGASLEPDPNLEVEVDLSKSQSTLRIKE-EDKTDATNISLLNVSVKLSISNSGNRVSQDQNLQYAREPLQHLEQLDQTQQEQQRLRRAYSLIPSWLRTLLGVLSLGQCRR
KC285152 -MAGTGASLEPDPNLEVEVDLSKSQSTLRIKE-EDKTDATNISLLNVSVKLSINNSGNRVSQDQNLQYAREPLQHLGQLGQTQQEQQRLRRAYSLIPSWLRTLLGVLSLGQCRR
```

**Genogroup Ic - California sea lion astrovirus etc**

```
                                                                                        ?????????????????
JN420351 MKSLLRSTRVGLAANPVDGRNLAAGLRTSRSQLIQSQRSRDELDHVVGLVNVSLNLSNVTLTNQVQQDQDLQFLSGPQQRLALWVRILLAILSLRCVC
JN420354 MKSLLRSTLVDLAANPVAGQSHAAGLRMSRSQLIQSQNRSDEPDHVVGLVNVSLRLSDVSLTVQESQDQGHLLVREQPPRLGLLARILLALLSLRCVC
JN420357 MKSLLRSTPVDLAVSPVDDHSHVVAPKMSRSQSIQNLKRSDEPDHVVGLVNVSLTLSNVSLTVQELQDQDLPLGREQRRPLALLARILLALLSLRCAC
JN420358 MKSLLRSTPVDLAVSPVDDHSHVVAPKMSRSQSIQNLKRSDEPDHVVGLVNVSLTLSNVSLTVQELQDQDLPLGREQRRPLALLARILLALLSLRCAC
JN420356 MKSLLRSTPVDLAANPVGGQSHVAALRMSRSQSILNQRSSDEVDHVVGLINVSLQLSNVNLTVQELQDPEPLLDRQLQPHWARLARILLAILSLKCVC
```

**Genogroup Ib - canine astrovirus etc**

```
                                                                                   ?????????????????
KX599351 MSPLRLKPPEQNQHPPGANPGGETGMSKSLSTHNQSQTRGDETDLTIVVARELSLSLNGNSIRQESQDQNQQSPKLLHLLWVLLARILRVRLS
KX599349 MLPLRLNPPGQSQHPPGVNPGDGTGASRSLSTHNQGQIEGDETDLTIVVARELRLSLNGSSIRLESQDQDQQSPRLLHLLWGLLARILRVR--
KX599353 MLPLRLNPPGQSQHPPGVNPGDGTGMSKSLSTHNQGQIEGDETDLTIVVARELRLSLNGSSIRLESQDQNQLSPRLLHLLWELLARILRVQ--
KX756441 MLPLRLNPPVQNQHPPGANPGAGTGMSKSLSIHNQEQTEGDETDLTIVVARELRLSLNGSSIKLELQDQGQQSPRLLHLLWELLARILRVR--
MF973501 MSPSRLNPPVQTITPPGANPGDVTGMLRSQSIPNHKK-DADETDLTIVVAREMSLSLNGNLIRQELQDQNQQSPKLLHLLWGLLARILRVRLI
KP404150 MLPLRLNPPGQNQHPPGANPGAETGMSKSLSTHNQGQTEGDETDLTIVVARELRLSLNDSSIKLESQDQGQRLPKLLHLLWGLLARILRAQ--
KP404149 MSPLRLKPPEQNQHPLGANPGGGTGMSRSQSIHNPKQIGGDETDLTIVVARELRLSLNDSSIKLESQDQGQRLPKLLHLLWELLARILRVQ--
MF973500 MLPSRLNPPVQIITPPGASPGDVTGMLRSQSIPNHKK-GADETDLTIVVARELSLSLNGNSIRQELQDQNQQSPRLLHLLWGLLARILQVRLN
KX599350 MLPLRLKPPGQSQHPPGVNPGDGTGASRSLSTHNQGQIEGDETDLTIVVARELRLSLNGSSIRLESQDQDQQSPRLLHLLWGLLARILRVR--
KX599352 MLPSRLNPPVQTTTPPGASPGGVTGMLKLQSIPNHKK-DADETDLTIVVAKELSLSLNGNSIRQELQDQNQQSPKLLHLLWGLLARILRVRLN
```

**Genogroup III - Marmot astrovirus**

```
         sssssssssssssssssssssssss
KY855439 MRRLPSPLRLLVVVLGGFLAGCAVGELDRRNNNQTYNQPLLQPLLVELLQYPMLLALVVGTVLGVVATLLVAQLTRQSRPPLERLVRTRETTLRQNL
KY855441 MRRSPSPLRLLAVVLGGFLAGCAVGELERRGNNQTHSQPLLQPLLVELLQYPMLLALVVGTVLGVVATLLVVQLTRQSRLPLERLVRTRETTLRQSL
KY855442 MRRSPSPLRLLAVVLGGFLAGCAVGELERRGNNQTHSQPLLQPLLVELLQYPMLLALVVGTVLGVVATLLVVQLTRQSRLPLERLVRTRETTLRQSL
KY855440 MRRSPSPLRLLAVVLGGFLAGCAVGELERRGNNQTHSQPLLQPLLVELLQYPMLLALVVGTVLGVVATLLVVQLTRQSRLPLERLVRTRETTLRQSL
```

**Genogroup IIId - porcine astrovirus 4 etc**

```
KY940075 ----MSNLKWSRLLRRCLTVVVGVAAGALIAYLQLLQPLLGELKLLNHLSGLVVGAIIGLAILISRLAPSSDRK----
LC201606 ----MSNPRWLRLRRRLLVVVVGVAAGAILAHLHLVRPLLGRWQFLDNLVGLLAGALIGLAILLNPAAPSSDKG----
LC201613 ----MSNPRWSRLLRRLLVVVVGVAAGGLLAHLRLVRPLLGGSPLLDNLVGLAVGAVIALALLVNRLLAASGKESQQH
KU764484 ----MSNPRWLRPLRRLLVVVVGAAAGAFLAHLHLIRPLLGRLPLLDNLVELLVGVAIALVNLAVRRLGALGKK----
KU764485 ----MSNPRWLRPLRRLLVVVVGAAAGAFLAHLHLIRPLLGRLPLLDNLVELLVGVAIALVNLAVRRLGALGKK----
KU764486 ----MSNLRWLRPLRRLLVVVVGVAAGALLAHLHLIRPLLGRLPLLDNLVEILVGVAIALVNLAVRHLGASDKK----
JF713713 ----MSSLRWSRLLRRCLTVVVGVAVGALIAYLQLLQPLLGELKLLSHLLGLAVGAIIGLAILINRVVPSSNKK----
LC201605 ----MSNPRWSRLLRRLLVVVVGAAAGVLLAHLRIIQPLLGKLPLLDNLLGLTVGAIIALAMLVNRLLPASDRE----
KX060809 ----MSNPRWSRLPRRLLVVVVGVAAGALLAHLQLTRPLLGRLPLLDNLVGLAVGAVIALALLVNRLLAASGNGSQQH
JQ340310 ----MSNPRWSRLRRRLLVVVVGVAAGALLAQLHLVRPLLGRWQFLDNLVGLLVAALIGLAILINPAAQPSNKE----
LC201609 ----MSNPRWLRLLRRLLVVVVGVAAGALLAHLRLARPLLGRLPLIDNLAGLVVGAVIALALVVNRLLEESDKGSQQH
KY933398 ----MSNPMWLRLLRRLLVVVVGVAAGGLLAHLRLVRPLLGRLPLIDNLVGLLVGAVLSLAILLSRVTRCTHKE----
LC201604 ----MSNLRWSRLLRRLLVVVVGVAAGALIAHLKLLQPLLGKLPLLDNLIGLVVGAVIALALLANRVLPSSNKG----
JX556692 ----MSNLRWSRLLRRCLTVVVGVAVGALIAYLQLLQPLLGELKLLNHLLGLAVGAIIGLAILINRVAPSSNKK----
LC201608 MTKTMSNPRWLRPLRRLLVVVVGVAAGALLAHLHLTKLLLGGSPLLDKLVGLIVGAVIALAILVGRHLGASGKE----
LC201603 ----MSNPLWLRLLRRFLTVVVGAAAGVLIAHLNLVQPLLGRSPLLDNLLGFAVGVVIGLAILTNKVLTSLSKK----
LC201600 ----MSNPRWSRLLRRLLVVVVGVAVGGILAHLRLVRPLLGGLPLLDNLVGLAVGAVIAMALLVNRLLVASDKGSQQH
LC201601 ----MSNPRWSRLLRRLLVVVVGVAVGGILAHLRLVRPLLGGLPLLDNLVGLAVGAVIAMALLVNRLLVASDKGSQQH
LC201602 ----MSNPRWSRLLRRLLVVVVGVAVGGILAHLRLVRPLLGGLPLLDNLVGLAVGAVIAMALLVNRVLPSSNKG----
KX033447 ----MSNPRWSRLRRRLLVVVVGVAAGALLAQLHLVRPLLGRWQFLDNLVGLLVVALIGLAILINPAAQHSGKQ----
KY214437 ----MSNPRWLRLLRRLLVVVVGVAVGGLLAHLRLVRPLLGRLPLIDNLAGLVVGAVIALALVVNRLLGVSGKGSQQH
KX060808 ----MSNLRWSRLLRRCLTVVVGVAAGVLIAHLNLVQPLLGKLPLLDNLVGLAVGAVIALVFLINRVLPSSSKE----
LC201610 ----MSNLKWSRLLRRCLTVVVGVAAGVLIAHLNLVQPLLGKLPLLDNLVGLAVGAVIALVLLINRVLPSSNKG----
LC201611 ----MSNLKWSRLLRRCLTVVVGVAAGVLIAHLNLVQPLLGKLPLLDNLVGLAVGAVIALVLLINRVLPSSNKG----
LC201612 ----MSNLKWSRLLRRCLTVVVGVAAGVLIAHLNLVQPLLGKLPLLDNLVGLAVGAVIALVLLINRVLPSSNKG----
```

**Genogroup IIIa - murine astrovirus etc**

```
         sssssssssssssssssssssssss
MF175075 MPRPSLLQLSLVAVVGGLAGALYGAALQALLTPQQRQQLFGLFFAAIPGLAVIAAGVGLLSGRLLARLSRRLRRPSERLARTRVIRSSLRWQRSSTRR
JX544743 MPRPSLLQLSLVAVVGGLAGALYGAALQALLTPQQRQQLFGLFFAALPGLAVIAAGVGMLSGRLLARLFRRLRRPSERLARTRAIRSSLRWQRSSTQR
JX544744 MPRPSLLQLSLVAVVGGLAGALYGAALQALLTPQQRQQLFGLFFAAIPGLAVIAAGVGMLSGRLLARLSRRLRRPSERLARTRAIRSSLRWQRSSAQR
KT946731 PRLPSPLSLAIVAV-GGVVGGAVLAQLDRLLQQQGQLPH--LVLAAVPGLAIGLAGATLVA-KLLPPSRKQSQQHWEPWDPIRAIP------------
KT946730 ----------MVAV-SGLTGGAALAALEKLLQPQGLPPF--LLLLAVPGLVVVGLSAGLVAER-LTRSTKPSRRHSVP--------------------
KT946728 ----------MVAV-SGLTGGAALAALEKLLQPQGLPPF--LLLLAVPGLVVVGLSAGLVAER-LTRSTKPSRRHSVP--------------------
KT946729 ----------MVAV-SGLTGGAALAALEKLLQPQGLPPF--LLLLAVPGLVVVGLSAGLVAER-LTRSTKPSRRHSVP--------------------
KT946726 ----------MVAV-SGLTGGAALAALEKLLQPQGLPPF--LLLLAVPGLVVVGLSAGLVAER-LTRSTKPSRRHSVP--------------------
KT946727 ----------MVAV-SGLTGGAALAALEKLLQPQGLPPF--LLLLAVPGLVVVGLSAGLVAER-LTRSTKPSRRHSVP--------------------
```

* KT946731 lacks the ORFX AUG codon - possibly a defective sequence

**Genogroup III - Rodent astrovirus**

```
         sssssssssssssssssssssssssssssss
KT946735 MPRLRSPLPRLVALVSGLAVGLWCAGIVKATGLPQQQQLFLHALATPGLVGFALALSVLVGMLQTPSPPRSPLHLVP
KT946736 MPRLRSPLPRLVALLSGLAVGLWCAGTVKATGLPQQQQLFLHALATPGLVGFALALSVLVGMLQTPSPPRSLLHSVP
```

**Genogroup III - Bovine astrovirus**

```
                                                             ????????????????????
LC047800 MAGRIGTLREWWSNLLQATNRHRPPPVGDVPDAELSMSTLIQ-DPHNSRRDSVPVQACACALTGVAAGVNKIRASFRYVRRSPQP---
KP264970 MVGRVGTPREWWSNLLQAINRHKRRRVRDANDGKLSMSMSTH--GLSHRTDSEQMLGCACALTGVLEGVKLPRVSLLSIRR-------
LC047798 MVGRTGTPHEWWSSLLQAINRQRRPRALDVRDGQQSMSMSTL--DLSQRRNSAPALACVCGLTAVVAGVNKVRASIKYIKR-------
LC047790 MAGRTATLRKWWCSLLQAINRL-PRCDLDAPDGPLSMSMSTQTEEILTETPMAPSLTCGCGLTAVFAGVNQIRASMKFTKPLQPPSAP
```

**Genogroup IIIb - bovine astrovirus etc**

```
LC047799 MDQLLHKVEQIPSSSSRG-RDEPDGGDSKIK---LGCLPFRLRLRVPGV-VSVR--VRALSTRRLSQHLAQWARIIRRQ-------
LC047789 MDQLLHKVEQIPNSSSRG-RDEPDGGDSKIK---LGCLPFRLRLRVPGV-VSVR--VRALSTRRLSQHLAQWARIIRRQ-------
LC047794 MDQLLHKVEQIPNSSSRG-RDEPDGGDSKIK---LGCLPFRLRLRVPGV-VSVR--VRALSTRRLSQHLAQWARIIRRQ-------
LC047795 MDQLLHKVEQAPNSSSRG-KDEPDGGDSKIR---LGCLPFKLRLRVPGV-VSVR--VRALSTRRLSQHLAQWARIIRRQ-------
LC047801 MDQLPHKVEQAPNSSSRG-KDEPDGGDSKIR---LGCLPFKLRLRVPGV-VSVR--VRALYTRRLSQHLAQWARTIRRQ-------
LC047788 MDQLPQQVGRPNSSSNRE-GDEPDGGDSRIR---LGCLPFRMQVRVPGV-VSVR--VRALCTSRLSRRLERWARIIQIK-------
LC047791 MDQPPHQVGRPNNSSSREEGDELDGGEGKLQ---LGCHPFRLRLRIPGV-ASVR--IRELSISGSSRRLGRLAQIILNKLKLN---
LC047792 MDQLPHQAGQPNSSSSREEGDEIDGGEGRFQ---LGCLPFRLRLRIPGV-ASVR--IRELSTSGLSQRSERLAQIILNRLKLN---
HQ916314 MDQLPPKVEAASQDSNN--SSVVDGAEQDHRPRSCGCFPTKLRVA-GGG-LGFHGSITVWWSRRLSQLSVRSAPMAQAQ-------
HQ916316 MDQLPPKVGAANQDNNS--SSAVDGAGQGRQPRWCGCYPSKIRVA-GGG-FGLRGSVTVWFTRRLSQLSAR--------------
MG660832 MEQLPTRQELSEDG-----VDGGTGETAKRRRSTCGCYPTKLRVS-AGI-LAARASVTVWFSRRSTQPSVQLDPTEVNRLNASLRA
KM822593 MDQLPHQVGQPNNNSSRGERDEIDGGEGKLQ---LGCLPFRLRLRVPGV-ASVR--IRELSTSELSQRLGRLARIIQTK-------
LC047797 MDQLPPKAEGSKLVCH---ADAVDGCKCKRCKH-RGCLPIG-RIRLSGAFLAARMSITVSFIRRSLRRSGP--------------
HQ916313 MEQLPTRQELPEDS-----VDGGAGGTTRRRRSTCGCYPTKLRFS-ARI-LAARASVTVWFSRRSTQPSVQLDPTEVSRLNASSRA
KJ620979 MEQLPTRQELSEDS-----VDGGTRGTTGRRRSTCGCYPTELRVS-AGV-LAARASVTVWFSRRSTQPSVQLDQTEVNRLNANSRA
KJ620980 MEQLPTRQELSEDS-----VDGGTRGTTGRRRSTCGCYPTKLRVS-AGV-LAARASVTVWFSRRSTQPSVQLDQTEVNRLNANSRA
HQ916317 MEQLPTRQELSEDS-----VDGGTGGTTKRRRSTCGCYPTKLRVS-AGI-LAARASVTVWFSRRSTQPSVQLDPTEVNRLNASSRA
LC047787 MEQLQPKVGSGKDSSKLH-ADDVDGCKCRRCRH-RGCLPVR-RIRLAGVFLAVRMSITASFTRRSRRRSGP--------------
LC047796 MEQLPTRQELLEDS-----ADGGAGGITRPRRSTCGCYPTKLRFS-AGF-LAARASVTVWFSRRSIQPSVQLDQTEVNRLNASLRA
```

**Genogroup IIIc - porcine astrovirus 2 etc**

```
                                                ????????????????????
LC201590 --MAPARIRRAHPGPSRQEGEET--GRNH---KSISGCWPTKLRVSAGFLAARVSITAWFSRRSTQHSAPSDQMEASRLSVS---------
KP747573 --MAPPRIRRSLPEQIRQEGEGT--GEDHS--RPWCGCYPTKVRVAGAILGLRASVTVWLSRRLSQLSGQSDPMGVDQSRQNSPSSSTPAQ
KY940077 --MAPARIRRALPELNRQGEEGT--GENR---RLTSGCLPIKIRLSAGVLAARVSVVVWFSRRSRRPLAQSDQMVVSKSSVK---------
KP982872 --MAPARIRRALPEPNRQEEEGT--GESHS--RPWCGCYPTKIRVAGAFLGLRASVTVWLSRRLSRLSEQLDPTARDRSRQSSLSCSTPAQ
LT898434 --MAPARIRRALPEHNRQGGEET--GGNR---RSTSGCWPVKIRLSAGILAARVSVVVWFSRRSRQPSAQSDQMVVTRSSVR---------
LC201588 --MAPARIRRAHPGLNRQDEGET--EENHS--KPSCGCYPTRIRVAGAILGLRASVTVWLSRRLSQPSGQSGPTELDRSRQSLQSYSIPAQ
KP759770 --MAPPRIRRALPEQIRQEGEGT--GENHN--RPWCGCYPTRFRVAGAILGFRASVTVWLSRRLSQLSGQSDPMGVGQSRQNSLSSSTPAQ
KY214438 --MAPPRIRRALPEQIGQEGGET--EGNHN--RPLCGCYPTRVRVAGAFLGLRASVTVWLSRGLSRLSARSGRMGQAK-------------
KR868724 --MAPPQIRRALPRQSRQEGGEV--EGNH---RLTSGCCPIKVRLSASFLAARASVIVWFSRKSIRPLEQLDQMAVNK-------------
KR868723 --MAPPRIRRALPRQDKQGEGET--GENR---RSISGCCPVKIRLSAGFLAAKVSVVVWFSRRSIRPLGQLDPMAVNKSSVR---------
KR868722 --MAPPRIRRALPRQDKQGEGET--GGNH---RLISGCCPIKIRLSARFLAARVSVVVWFSRRSTRPLGQLGPMAVSKSSVR---------
KR868721 --MAPPRIRRALPRQDKQGEGGT--EGNH---RLISGCCPIKIRLSARFLAARVSVVVWFSRRSTRPLGQLGPMAVSKSSVR---------
LC201593 --MAPPRIRRALPEPNRQGDEGT--REDHS--QSTCGCLPTKVRVAGAVLGFRASVTVWWHRGLSRLSGRSVPTDPGT-------------
KY940076 --MAPARIRRALPEPNKQGEEGTEQGDRP---GSTCGCFPIKVRVAAGFLGARVSVIAWFSRRSTPRLAQLGRMRAERLSVN---------
LC201592 --MAQPRIRRALPALGRDAGGGT--GENNNNQRSTCGCYPTKIRVAGAFLGLRASVTVWWSRGLSRLSGRLGPTGLEK-------------
LC201587 --MAPARIRRARPGPSRQEGEEV--GENHS--RPWCGCYPTRIRVAGAILGLRASVTVWLSRRLSRPSEQSGPMEPDRSRQNSLSCSIPVQ
LC201589 MEQLPPR-RGRLVRRQAIEEGGT--GENP---KSTSRCWPTKVRLSAACLAAKGSITVWFSRRSTQHLERSAQTEVERSSVS---------
LC201594 MDRLPPR-RGRLARQQATEEGETE-GDRS---RPLCGCYPTKIRVAGAVLGARASVTVWLSRRLSQLSAQLGPMVADQSRRSSPSS-----
LC201585 MEQLPPR-RGRLARQQA-EGGETEQGDSP---RSMCGCYPLKLRVAAGFLGARVSVVAWFSRRSTRRSAQSARMGVNK-------------
LC201586 MEQLPTR-RGRLARQQATEGEGIEQGDRP---RSTCGCFPVKIRVAAGFLGARVSVVALFSRRSTRLLAQSDPMGVNK-------------
KJ495986 MEQLPPR-RGRLARQQ-VGGEETEQGDNP---KSTCGCYPLKLRVAAGFLGARVSVVAWFSRRSTQRSAQSAQMVVTK-------------
JF713710 MEQLPPR-RGRLARQQAVGGEEIEQGDNP---RSTCGCYPLKLRVAAGFLGARVSVVAWFSRRSIQHSAQLVQMEVSK-------------
JF713712 MEQLPPR-RGRLERQQATEGGGIEQGDNH---NSTCGCFPVKIRVAAGFLGARVSVVALFSRRLTQPSARLDPTGVSK-------------
JX556690 MEQLPPR-RGRLARQQAIGEGETEQGDNH---RSMCGCYPLKLRVAAGFLGARVSVVAWFSRRSIQHSAQSVQTEVSK-------------
```

**Supplementary Figure 9 | Sequences of putative XP proteins encoded in diverse astroviruses.** Amino acid alignments of putative XP proteins in different astrovirus groups. Sequences were aligned with MUSCLE [1]. Sequences are listed in the same order as in the ORF1b phylogenetic tree (Fig. S1). Amino acids are colour-coded according to their physicochemical properties. Transmembrane regions predicted by Phobius [5] are indicated with pink bars above the alignment. Potential transmembrane regions (hydrophobic regions scored below threshold by Phobius) are indicated with pink question marks. Signal peptides predicted by Phobius are indicated with pink “s”s. In all cases, predictions are based on the first sequence in each alignment.

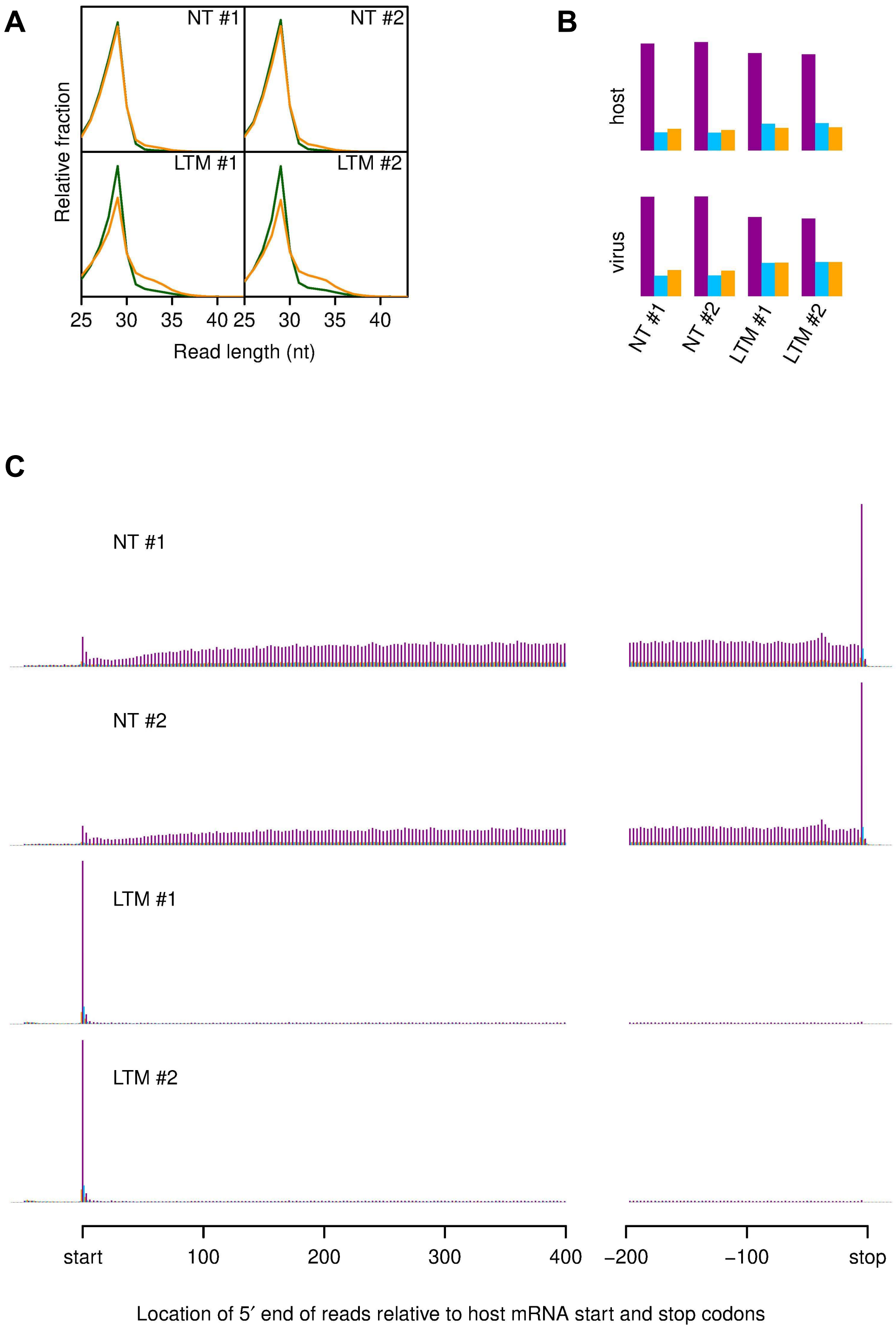


**Supplementary Figure 10 │ Assessment of ribosome profiling quality.** Cells were harvested at 12 hpi and either flash frozen with no pre-treatment (NT), or pre-treated with lactimidomycin for 30 minutes followed by flash freezing (LTM). **(A)** Relative length distributions for Ribo-Seq reads mapping to virus (orange) and host (green) mRNA coding regions. NT - no **(B)** Phasing of 5′ ends of RPFs (≥25 nt) that map to the viral ORFs (excluding dual coding regions) or host mRNA coding regions. **(C)** Histograms of approximate P-site positions of RPFs (≥25 nt) relative to annotated initiation and termination sites summed over all host mRNAs.

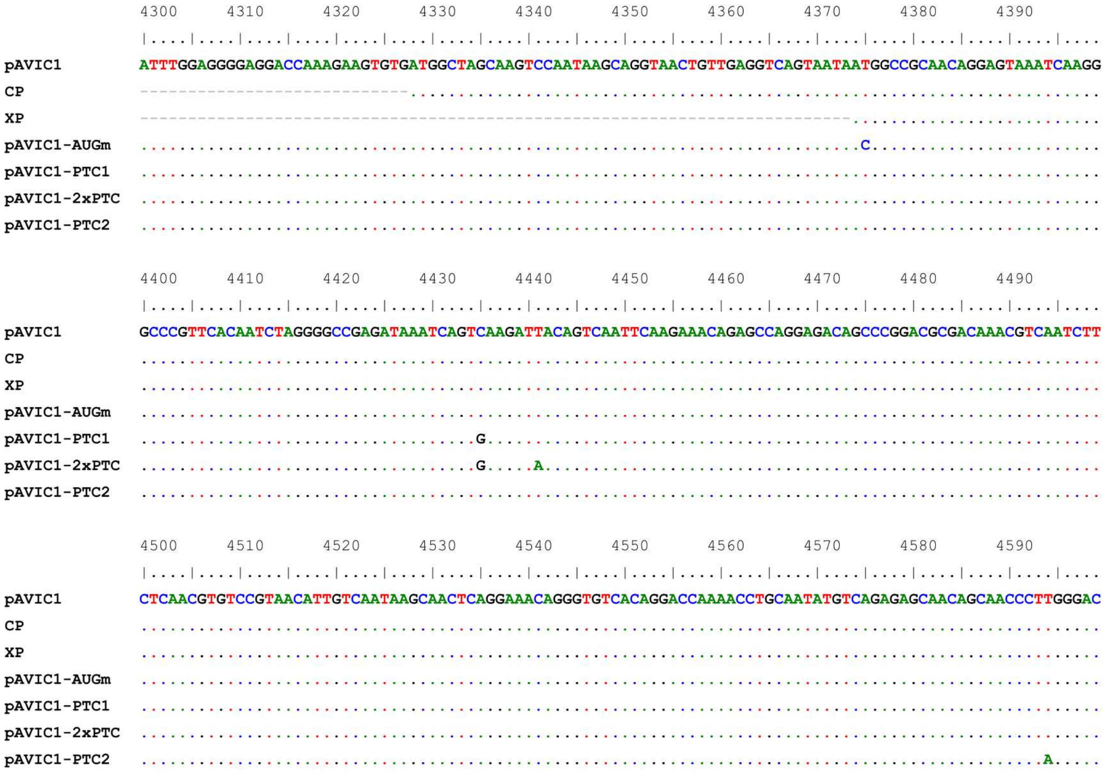


**Supplementary Figure 11 │ Nucleotide alignments of pAVIC1 mutants.** Alignment of pAVIC1, CP (ORF2), XP (ORFX), pAVIC1-AUGm, pAVIC1-PTC, pAVIC1-2×PTC and pAVIC1-PTC2 mutants. Only nucleotide differences from wt pAVIC1 are shown for the mutants.

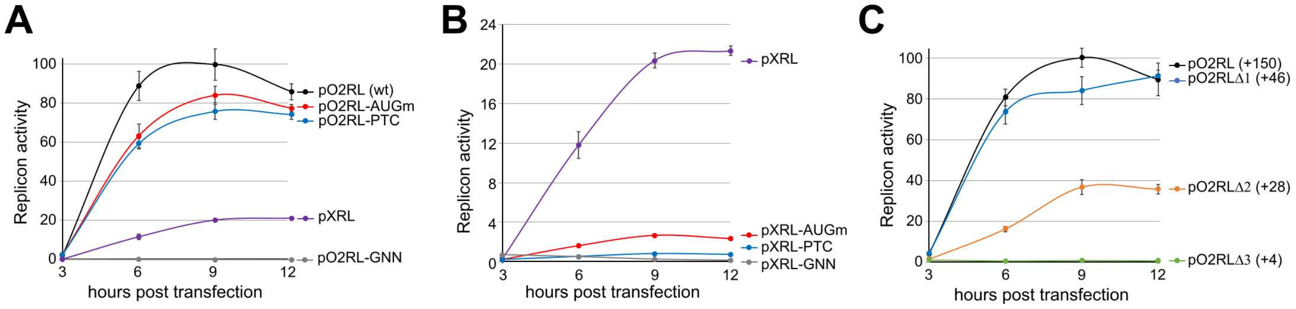


**Supplementary Figure 12 │Replicon activity in Huh7.5.1 cells.** Relative replicon luciferase activities representing translated product associated with activity of the subgenomic promoter, measured after RNA transfection of Huh7.5.1 cells (mean ± s.d.; *n* = 3 biologically independent experiments). Values are normalized so that the maximum wt value is 100%.

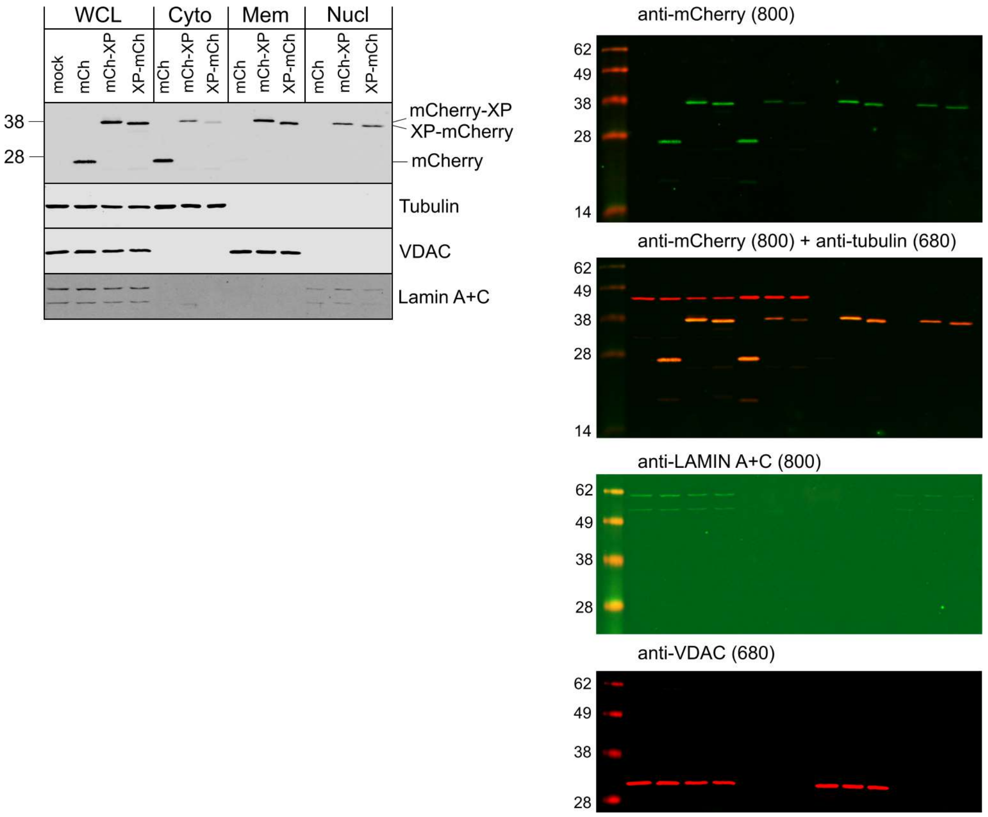


**Supplementary Figure 13 | Original western blot scans for Fig. 5B.**

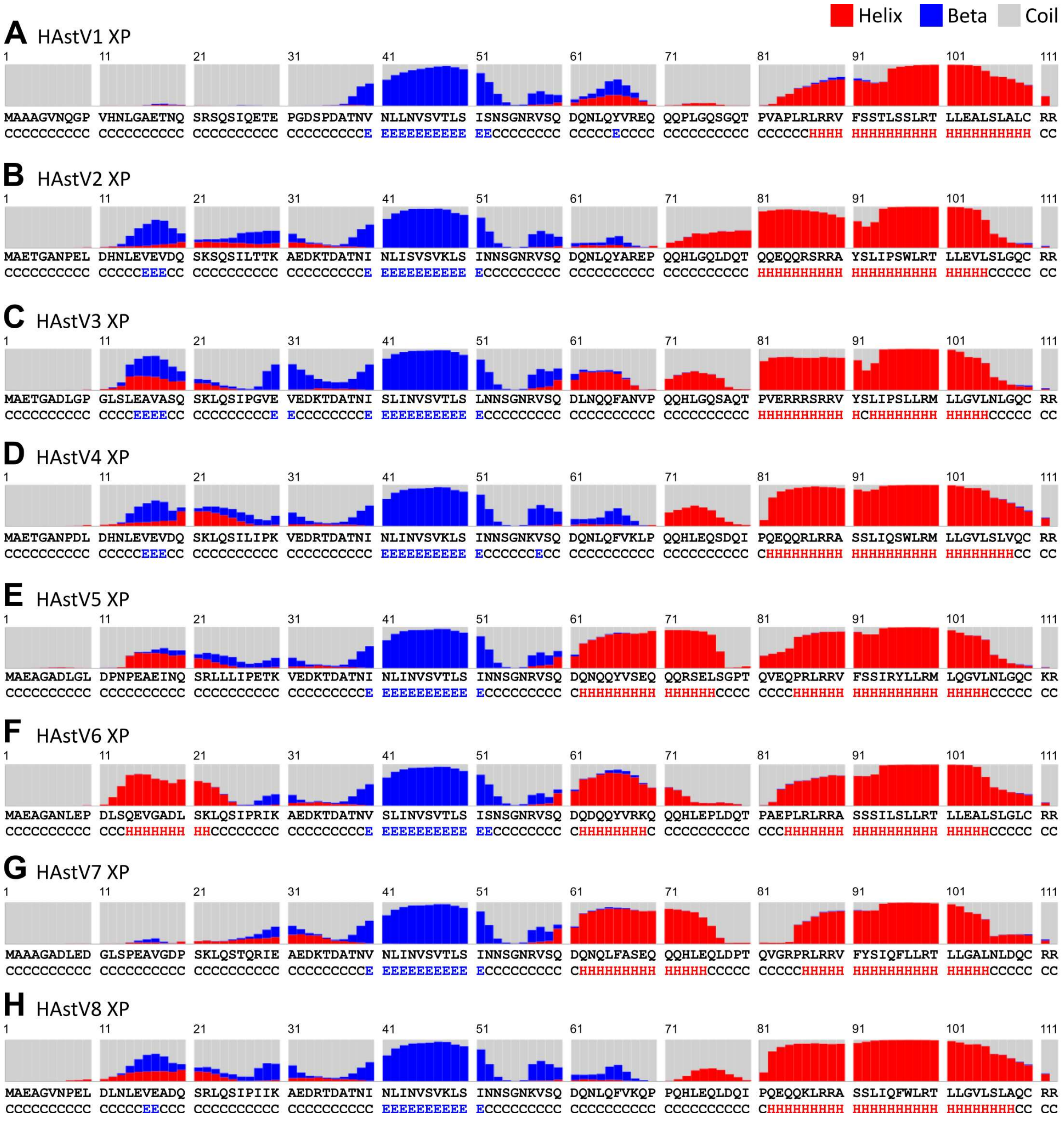


**Supplementary Figure 14 │ Predicted secondary structure of HAstV XPs.** Protein secondary structures for representative sequences (see Fig. 1F) were predicted with RaptorX [6].

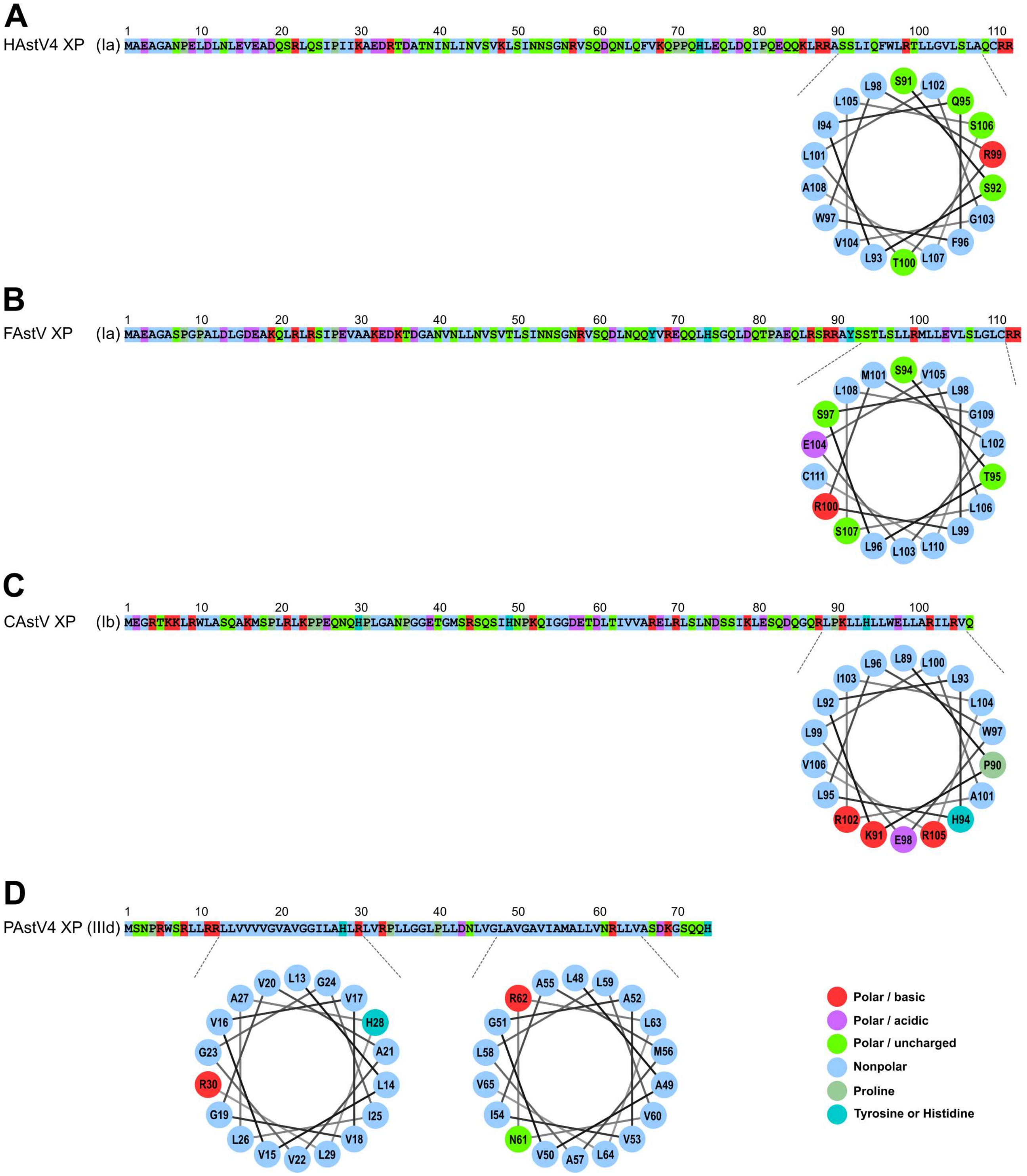


**Supplementary Figure 15 │Helical wheel representations for TM regions of other astrovirus XPs used in Fig. 6I.** TMs were predicted with Phobius [5] (http://phobius.sbc.su.se/cgi-bin/predict. pl) and helical wheels were created and analysed using Heliquest [7] (http://heliquest.ipmc.cnrs.fr).

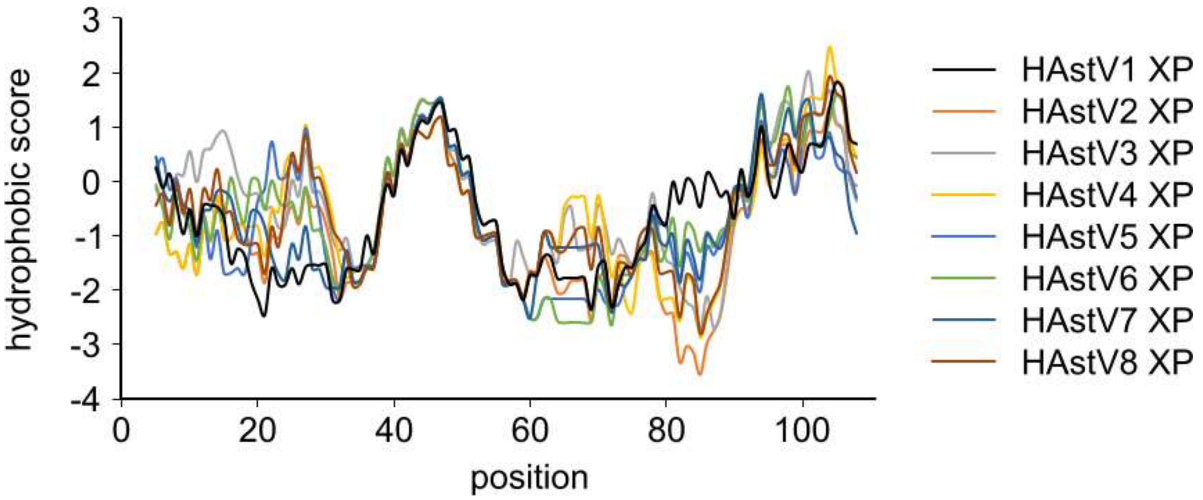


**Supplementary Figure 16 │Individual Kyte-Doolittle hydropathy plots for HAstV1-8 XPs.** Hydropathy plots for HAstV1 pAVIC1 and representative HAstV2–8 sequences (see Fig. 1F) were predicted with protscale (https://web.expasy.org/protscale).

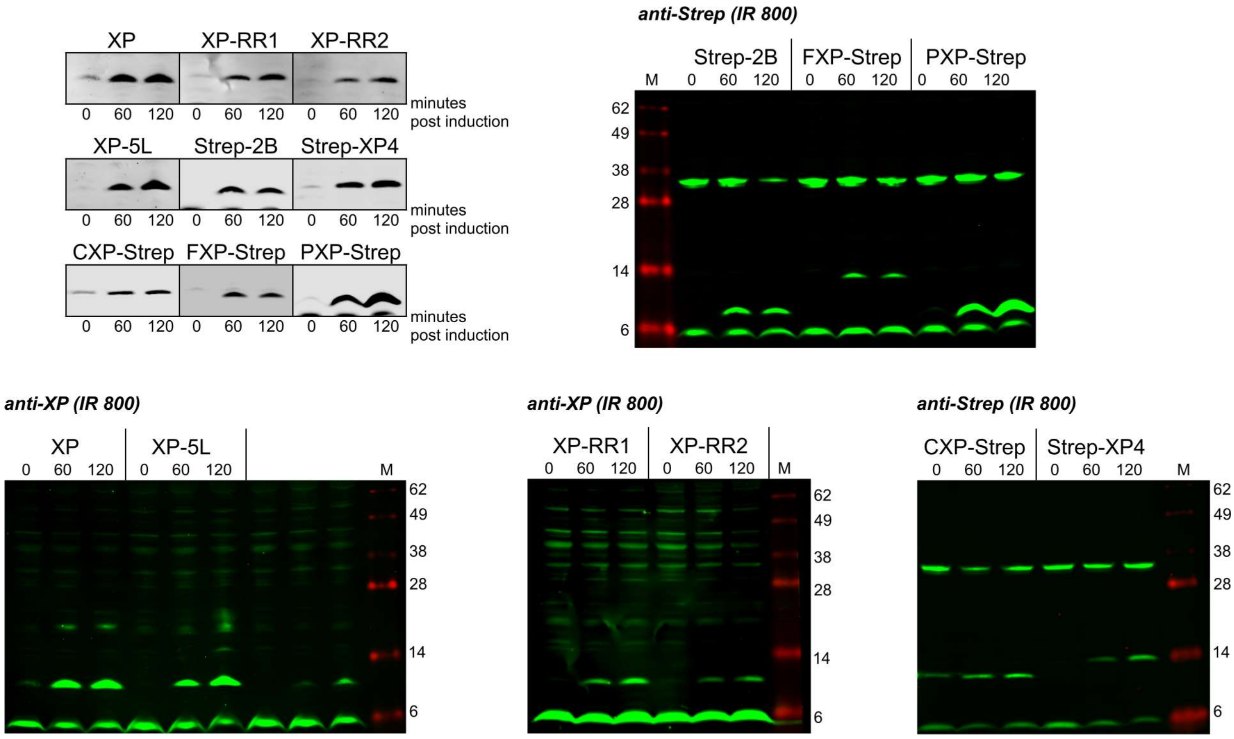


**Supplementary Figure 17 | Original western blots for Fig. 6J.**

**Supplementary Table 1 │ XP and YP statistics in different astrovirus groups**

| group | protein | *n* | median mass (kDa) | median pI | median length (aa) | predicted TM(s) |
|---|---|---|---|---|---|---|
| Bat astrovirus | XP | 1 | 17.1 | 13.4 | 148 | yes |
| Genogroup IId - bovine astrovirus etc | XP | 17 | 10.7 | 12.7 | 91 | yes |
| Genogroup IV - MLB astroviruses | XP | 13 | 8.4 | 8.6 | 78 | yes |
| Rodent astrovirus | XP | 4 | 8.3 | 5.5 | 79 | yes |
| Genogroup Ia - human astrovirus etc | XP | 39 | 12.3 | 8.0 | 112 | no |
| Genogroup Ic - California sea lion astrovirus etc | XP | 5 | 10.8 | 7.9 | 98 | no |
| Genogroup Ib - canine astrovirus etc | XP | 10 | 10.1 | 9.7 | 91 | no |
| Marmot astrovirus | XP | 4 | 10.7 | 12.1 | 97 | yes |
| Genogroup IIId - porcine astrovirus 4 etc | XP | 25 | 7.6 | 12.5 | 70 | yes |
| Genogroup IIIa - murine astrovirus etc | XP | 8 | 7.5 | 12.1 | 73 | yes |
| Rodent astrovirus | XP | 2 | 8.0 | 12.5 | 77 | yes |
| Bovine astrovirus | XP | 4 | 9.1 | 11.5 | 82 | no |
| Genogroup IIIb - bovine astrovirus etc | XP | 18 | 8.2 | 11.4 | 72 | no |
| Genogroup IIIc - porcine astrovirus 2 etc | XP | 24 | 8.3 | 11.8 | 75 | no |
| Genogroup IIc mamastrovirus 10 etc | YP | 6 | 9.9 | 6.2 | 91 | no |

**Supplementary Table 2 │ Host and virus read counts for different Ribo-Seq samples**

| drug | repeat | time point | total reads | host rRNA | host mRNA | vRNA |
|---|---|---|---|---|---|---|
| No treatment | NT #1 | 12 hpi | 34,261,071 | 23,068,841 | 3,960,680 | 778,926 |
| | NT #2 | 12 hpi | 41,436,501 | 27,367,513 | 5,669,906 | 1,035,209 |
| Lactimidomycin | LTM #1 | 12 hpi | 11,621,251 | 6,024,566 | 625,793 | 177,015 |
| | LTM #2 | 12 hpi | 12,903,118 | 9,177,384 | 828,982 | 240,408 |

**Supplementary Dataset 1 | Accession numbers of the 415 astrovirus ORF2 sequences**

AB000283 AB000284 AB000285 AB000286 AB000287 AB000288 AB000289 AB000290 AB000291 AB000292 AB000293 AB000294 AB000295 AB000296 AB000297 AB000298 AB000299 AB000300 AB000301 AB009984 AB009985 AB013618 AB025801 AB025802 AB025803 AB025804 AB025805 AB025806 AB025807 AB025808 AB025809 AB025810 AB025811 AB025812 AB031030 AB031031 AB037272 AB037273 AB037274 AB290149 AB308374 AB496913 AB823731 AB823732 AB829252 AB914705 AB914706 AF056197 AF117209 AF141381 AF248738 AF260508 AY179509 AY720891 AY720892 DQ028633 DQ070852 DQ344027 DQ630763 EF138823 EF138824 EF138825 EF138826 EF138827 EF138828 EF138829 EF138830 EF138831 EF583300 EU847144 EU847145 EU847155 FJ222451 FJ375759 FJ402983 FJ571065 FJ571066 FJ571067 FJ571068 FJ571070 FJ571071 FJ571072 FJ571073 FJ571074 FJ755402 FJ755403 FJ755404 FJ755405 FJ792842 FJ890351 FJ890352 FJ890355 FJ973620 FM213330 FM213331 FM213332 GQ267696 GQ405855 GQ405856 GQ405857 GQ415660 GQ415661 GQ415662 GQ495608 GQ502193 GQ891990 GQ901902 GQ914773 GU223905 GU376736 GU562296 GU732187 GU985458 HM045005 HM237363 HM447045 HM447046 HM450380 HM450381 HM450382 HM756258 HM756259 HM756260 HM756261 HQ398856 HQ623147 HQ623148 HQ647383 HQ668129 HQ668143 HQ916313 HQ916314 HQ916315 HQ916316 HQ916317 JF327666 JF491430 JF713710 JF713711 JF713712 JF713713 JF729316 JF742759 JF742760 JF755422 JN052023 JN088537 JN193534 JN420351 JN420352 JN420353 JN420354 JN420355 JN420356 JN420357 JN420358 JN420359 JN592482 JN887820 JQ081297 JQ086552 JQ340310 JQ403108 JQ408745 JX087963 JX087964 JX087965 JX544743 JX544744 JX544745 JX544746 JX556690 JX556691 JX556692 JX556693 JX684071 JX684072 JX857868 JX857869 JX857870 KC285152 KC342249 KC609001 KC692365 KC915034 KC915035 KF039910 KF039911 KF039912 KF039913 KF157967 KF211475 KF233994 KF374704 KF417713 KF499111 KF668570 KF787112 KF859964 KJ476832 KJ476833 KJ476834 KJ476835 KJ476836 KJ476837 KJ476838 KJ495986 KJ495987 KJ495991 KJ495992 KJ495993 KJ495994 KJ495996 KJ495999 KJ496001 KJ571486 KJ620979 KJ620980 KJ656124 KJ920196 KJ920197 KM017741 KM017742 KM017743 KM035759 KM358468 KM401565 KM822593 KP264970 KP404149 KP404150 KP404151 KP404152 KP663426 KP747573 KP747574 KP759770 KP862744 KP942582 KP942583 KP942584 KP942585 KP942586 KP942587 KP942588 KP942589 KP942590 KP942591 KP942592 KP942593 KP982872 KR349488 KR349489 KR349490 KR349491 KR868721 KR868722 KR868723 KR868724 KT224358 KT946726 KT946727 KT946728 KT946729 KT946730 KT946731 KT946732 KT946733 KT946734 KT946735 KT946736 KT956903 KT963069 KT963070 KT963071 KU764484 KU764485 KU764486 KX022687 KX033447 KX060808 KX060809 KX266901 KX266902 KX266903 KX266904 KX266905 KX266906 KX266907 KX266908 KX273058 KX599349 KX599350 KX599351 KX599352 KX599353 KX599354 KX645667 KX683863 KX756441 KY024237 KY024238 KY073229 KY073230 KY073231 KY073232 KY073233 KY214437 KY214438 KY271946 KY412124 KY412125 KY412126 KY412127 KY744137 KY744138 KY744139 KY744140 KY744141 KY855437 KY855438 KY855439 KY855440 KY855441 KY855442 KY859988 KY933398 KY933399 KY933670 KY940075 KY940076 KY940077 KY940545 L06802 L13745 L23513 LC047787 LC047788 LC047789 LC047790 LC047791 LC047792 LC047793 LC047794 LC047795 LC047796 LC047797 LC047798 LC047799 LC047800 LC047801 LC064152 LC201585 LC201586 LC201587 LC201588 LC201589 LC201590 LC201592 LC201593 LC201594 LC201595 LC201596 LC201597 LC201598 LC201599 LC201600 LC201601 LC201602 LC201603 LC201604 LC201605 LC201606 LC201608 LC201609 LC201610 LC201611 LC201612 LC201613 LC201615 LC201616 LC201617 LC201618 LC201619 LC201620 LC341267 LN879482 LT706530 LT706531 LT898424 LT898434 MF033385 MF033386 MF175073 MF175075 MF684776 MF973495 MF973496 MF973497 MF973498 MF973499 MF973500 MF973501 MG571777 MG660832 MG693176 S68561 U15136 Y08632 Y15937 Y15938 Z25771 Z33883 Z46658 Z66541
